# Broad phylogenetic and functional diversity among mixotrophic consumers of *Prochlorococcus*

**DOI:** 10.1101/2021.08.18.456384

**Authors:** Qian Li, Kyle F. Edwards, Christopher R. Schvarcz, Grieg F. Steward

## Abstract

Small eukaryotic phytoplankton are major contributors to global primary production and marine biogeochemical cycles. Many taxa are thought to be mixotrophic, but quantitative studies of phagotrophy exist for very few. In addition, little is known about consumers of *Prochlorococcus*, the abundant cyanobacterium at the base of oligotrophic ocean food webs. Here we describe thirty-nine new phytoplankton isolates from the North Pacific Subtropical Gyre (Station ALOHA), all flagellates ∼2–5 um diameter, and we quantify their ability to graze *Prochlorococcus*. The mixotrophs are from diverse classes (dictyochophytes, haptophytes, chrysophytes, bolidophytes, a dinoflagellate, and a chlorarachniophyte), many from previously uncultured clades. Grazing ability varied substantially, with specific clearance rate (volume cleared per body volume) varying over ten-fold across isolates and six-fold across genera. Slower grazers tend to create more biovolume per prey biovolume consumed. Using qPCR we found that the haptophyte *Chrysochromulina* was most abundant among the isolated mixotrophs at Station ALOHA, with 76–250 cells mL^-1^ across depths in the upper euphotic zone. Our results show that within a single ecosystem the phototrophs that ingest bacteria come from many branches of the eukaryotic tree, and are functionally diverse, indicating a broad range of strategies along the spectrum from phototrophy to phagotrophy.

## Introduction

Small eukaryotic phytoplankton in the ‘pico’ (< 3µm) and ‘nano’ (2–20 µm) size classes are often dominant contributors to phytoplankton biomass and primary production in open oceans [1][2][3] and are major drivers of nutrient cycling [4][5]. At the same time, many flagellate and ciliate taxa are known to be mixotrophic, capable of obtaining nutrition through combined photosynthesis (autotrophy) and phagocytosis (heterotrophy) [6][7]. Mixotrophs have been observed in sunlit habitats throughout the ocean [8][9] and are estimated to contribute about two thirds of total bacterivory in open-ocean Atlantic ecosystems [10]. Phototrophic and heterotrophic prokaryotes are themselves major contributors to pelagic biomass and production [11][12][13], and therefore the bacterial grazers that also photosynthesize may play a key role in regulating productivity, element cycling, and food web dynamics. To understand how these systems function it is necessary to quantify fundamental parameters such as grazing kinetics and growth efficiencies of the main bacterial grazers, and it may be particularly consequential if the main grazers are mixotrophs, because models predict that mixotrophic consumers increase primary production and carbon export, and decrease nutrient remineralization, relative to heterotrophic consumers [14][15].

Although the aggregate importance of pigmented flagellates for bacterial grazing has been documented [16][17], much less is known about which taxa are the major grazers, and how the broad phylogenetic diversity among these organisms translates into diverse ecological roles [8][18]. Progress in this area has been impeded in part by a paucity of representative cultured mixotrophs relative to the complexity found in natural communities [18][19], which often contain haptophytes, chrysophytes, dictyochophytes, chlorophytes, bolidophytes, cryptophytes, and dinoflagellates [20].

The North Pacific Subtropical Gyre (NPSG) is a chronically oligotrophic environment, where prokaryotic and eukaryotic phototrophs are dominated by *Prochlorococcus* [21] and small flagellates [22], respectively. Some of the likely grazers of *Prochlorococcus* in this habitat were identified by stable isotope probing by addition of ^13^C- and ^15^N-labeled *Prochlorococcus* MED4 cells to natural communities [23]. The 18S rRNA of various haptophytes, dictyochophytes, bolidophytes and dinophytes became significantly labeled, but most of the specific phylotypes identified in that study have not been isolated, and controlled lab studies of their grazing capabilities have been lacking. There has thus been no confirmation that most of these organisms directly ingest *Prochlorococcus* and no quantitative assessment of how ingestion rates and functional responses vary among them.

One exception is a recent study of the grazing ecophysiology of a phagotrophic mixotroph *Florenciella* (strain UHM3021; class Dictyochophyceae) isolated from the NPSG [24]. Given sufficient light, rapid growth of *Florenciella* was sustained by feeding on bacteria (*Prochlorococcus, Synechococcus,* or a heterotrophic bacterium) as the primary nutrient source, suggesting mixotrophy is an effective strategy for nutrient acquisition. The rate at which prey were ingested by this *Florenciella* strain was relatively low compared to heterotrophic flagellates of similar size, and prey ingestion was suppressed by high concentrations of dissolved nutrients. This was the first detailed characterization of *Prochlorococcus* consumption by a mixotrophic flagellate, and it remains unclear whether other small, open-ocean mixotrophs possess similar physiology. For example, *Florenciella* may be a relatively autotrophic mixotroph that relies on prey primarily for limiting nutrients [25]; other taxa may be more voracious grazers that rely on prey as a major energy source [26][27].

To broaden our understanding of how mixotrophs feed on one of the most important primary producers in the ocean, we isolated diverse mixotrophs (dictyochophytes, haptophytes, chrysophytes, bolidophytes, a dinoflagellate, and a chlorarachniophyte) from Station ALOHA, an open ocean site in the NPSG, 100 km north of the island of Oʻahu. By characterizing their grazing capabilities using *Prochlorococcus* as prey, and by quantifying their in situ abundances, we reveal functional diversity among the mixotrophs in this ecosystem and their contributions to *Prochlorococcus* mortality in situ.

## Methodology

### Isolation and cultivation

In total, 39 mixotrophic flagellates were investigated (Table 1). Thirty-three isolates were enriched and isolated from euphotic zone samples at Station ALOHA (22° 45’ N, 158° 00’ W) in February and May 2019. To select for mixotrophic grazers of *Prochlorococcus*, whole seawater was amended with K medium [28] (1/20 final concentration), and live *Procholorococcus* (MIT9301) was added as prey (∼5×10^6^ cells mL^-1^ final concentration). Enriched seawater samples were incubated under ∼70 µM photons m^-2^ s^-1^ irradiance on a 12 h:12 h light:dark cycle and monitored by microscopy daily up to five days. Each day, samples were serially diluted to extinction (9 dilution steps, 12 replicates per dilution) in 96-well plates in nutrient-reduced K medium (1/20 concentration) with a constant background of *Prochlorococcus* cells. Wells at the highest dilution showing growth of putative grazers were subjected to 3–6 further rounds of dilution to extinction. Four additional mixotrophs were isolated in full K medium using water from earlier cruises, and two (dictyochophyte strains UHM3021 described in [24] and UHM3050) were enriched in K minus nitrogen medium (K-N) without *Prochlorococcus* enrichment. All isolates were rendered unialgal, but not axenic, and maintained at 24 °C in K-N medium (∼0.2 µM N) amended with *Prochlorococcus* prey, under the same light conditions as above. *Prochlorococcus* fed to the cultures was grown in Pro99 medium [29]. Dense cells in late exponential-early stationary phase were harvested and concentrated through gentle centrifugation at 2,000 RCF for 5 minutes and resuspended in fresh K-N medium.

**Table 1.**
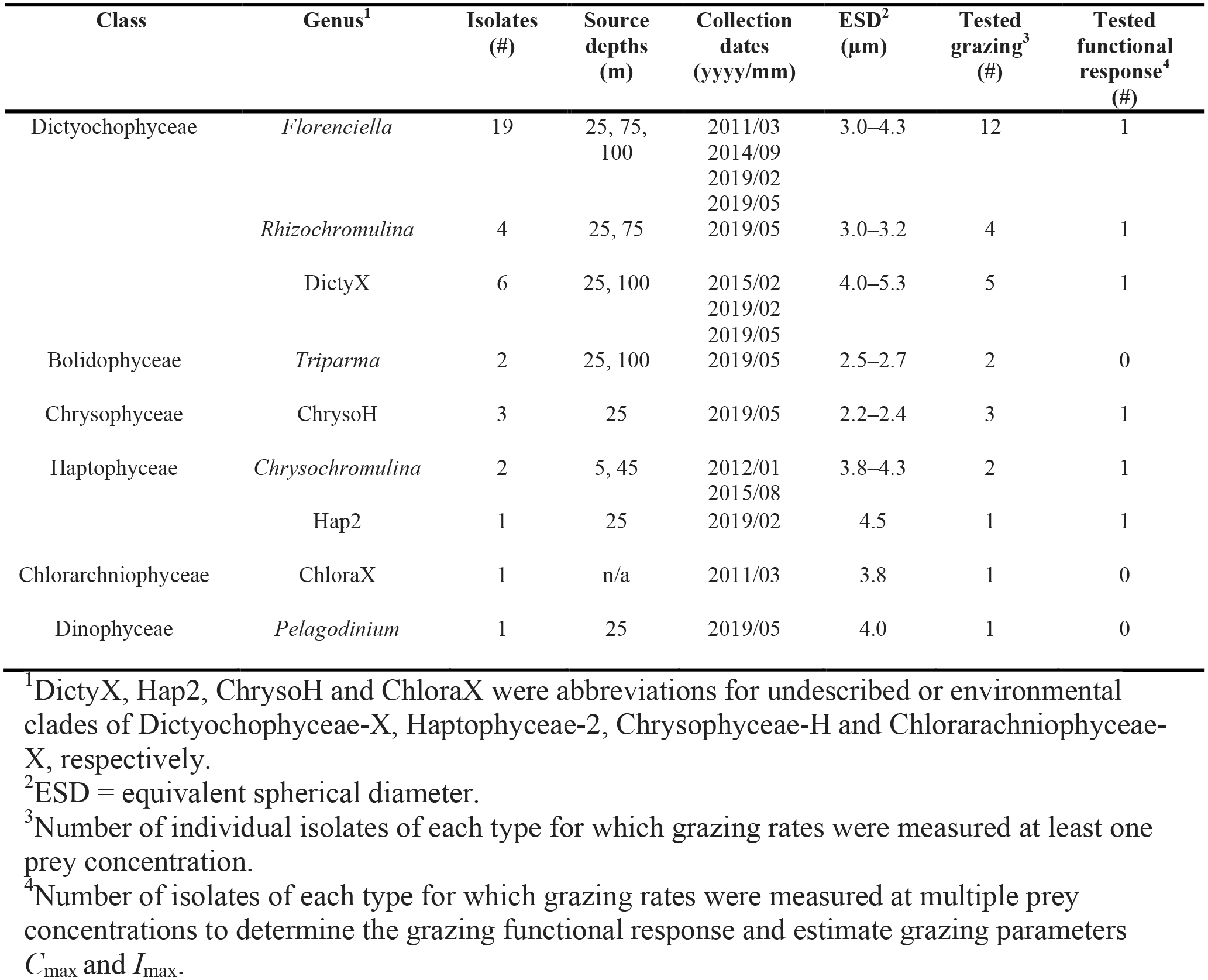
Mixotrophic flagellates surveyed in this study.

| Class | Genus<sup>1</sup> | Isolates (#) | Source depths (m) | Collection dates (yyyy/mm) | ESD<sup>2</sup> (μm) | Tested grazing<sup>3</sup> (#) | Tested functional response<sup>4</sup> (#) |
| --- | --- | --- | --- | --- | --- | --- | --- |
| Dictyochophyceae | *Florenciella* | 19 | 25, 75, 100 | 2011/03<br>2014/09<br>2019/02<br>2019/05 | 3.0–4.3 | 12 | 1 |
|  | *Rhizochromulina* | 4 | 25, 75 | 2019/05 | 3.0–3.2 | 4 | 1 |
|  | DictyX | 6 | 25, 100 | 2015/02<br>2019/02<br>2019/05 | 4.0–5.3 | 5 | 1 |
| Bolidophyceae | *Triparma* | 2 | 25, 100 | 2019/05 | 2.5–2.7 | 2 | 0 |
| Chrysophyceae | ChrysoH | 3 | 25 | 2019/05 | 2.2–2.4 | 3 | 1 |
| Haptophyceae | *Chrysochromulina* | 2 | 5, 45 | 2012/01<br>2015/08 | 3.8–4.3 | 2 | 1 |
|  | Hap2 | 1 | 25 | 2019/02 | 4.5 | 1 | 1 |
| Chlorarchniophyceae | ChloraX | 1 | n/a | 2011/03 | 3.8 | 1 | 0 |
| Dinophyceae | *Pelagodinium* | 1 | 25 | 2019/05 | 4.0 | 1 | 0 |

<sup>1</sup>DictyX, Hap2, ChrysoH and ChloraX were abbreviations for undescribed or environmental clades of Dictyochophyceae-X, Haptophyceae-2, Chrysophyceae-H and Chlorarchniophyceae-X, respectively.
<sup>2</sup>ESD = equivalent spherical diameter.
<sup>3</sup>Number of individual isolates of each type for which grazing rates were measured at least one prey concentration.
<sup>4</sup>Number of isolates of each type for which grazing rates were measured at multiple prey concentrations to determine the grazing functional response and estimate grazing parameters $C_{\max}$ and $I_{\max}$ .

### 18S rDNA sequencing and phylogenetic analysis

Cells were harvested by centrifuging 25–50 mL dense cultures at 3,000 RCF for 10 min at 4 °C. Genomic DNA was extracted from the pellets using the ZymoBIOMICS DNA Kit (Zymo Research, Irvine, CA, USA). A near-full-length section of the eukaryotic small-subunit ribosomal RNA gene (18S rDNA) was amplified by PCR with the Roche Expand^TM^ High Fidelity PCR System (Sigma-Aldrich, St. Louis, MO, USA) using either forward primer 5’- ACCTGGTTGATCCTGCCAG-3’ and reverse primer 5’-TGATCCTTCYGCAGGTTCAC-3’ [30], or Euk63F 5’-ACGCTTGTCTCAAAGATTA-3’ and Euk1818R 5’- ACGGAAACCTTGTTACGA-3’ [31]. Amplicons were purified using spin columns (DNA Clean & Concentrator-25; Zymo Research, Irvine, CA, USA) and sequenced (Sanger) using the same PCR amplification primers and an additional reverse primer 1174R, 5’- CCCGTGTTGAGTCAAA-3’ [32], when necessary to connect two ends. For phylogenetic analyses, similar sequences were retrieved from the PR^2^ database [33] based on BLAST similarity, and two environmental homologs (GenBank Acc. FJ537342 and FJ537336) were retrieved from NCBI GenBank for the undescribed haptophyte taxon, which was not affiliated with any reference sequence from the PR^2^ database. Sequence alignments including 39 isolates, 29 reference and 2 outgroup taxa were created with MAFFT v7.450 using the G-INS-i algorithm [34] in Geneious R11.1.5 (http://www.geneious.com) [35]. Terminal sites that lacked data for any of the sequences were trimmed and any sites with greater than 25% gaps were removed from the alignment, which generated a total sequence length of 1617 bp. Phylogenetic analysis was performed using MrBayes v3.2.6 in Geneious R11.1.5 [36] with two runs of four chains for 1,000,000 generations, subsampling every 200 generations with burn-in length 100,000, under the GTR substitution model. The Bayesian majority consensus tree was further edited within iTOL v5 [37]. All 18S rDNA sequences were deposited in GenBank with accession numbers MZ611704–MZ611740; MN615710–MN615711.

### Microscopic observation

The average diameter of flagellates in the exponential growth phase (n = 20 cells per strain) was measured by transmitted light microscopy using image analysis software (NIS-Elements AR, Nikon, Minato City, Tokyo, Japan) calibrated with a stage micrometer. Equivalent spherical diameter (ESD) and biovolumes were calculated assuming spherical cells. Chloroplasts were visualized by autofluorescence under epifluorescence microscopy. An average ESD of 0.64 µm was used for *Prochlorococcus* prey [24].

Visual evidence of phagocytosis was obtained by adding fluorescent beads (0.5 µm YG Fluoresbrite Microspheres; Polysciences) to each culture. Samples post incubation (∼2 h) were fixed with an equal volume of 4% ice-cold glutaraldehyde, and subsamples (20 µL) were mounted on a glass slide under a coverslip. Paired images captured using epifluorescence and transmitted light microscopy (Olympus BX51 with Leica DFC 7000 T color digital camera) were overlain to identify cells with ingested beads.

### Grazing experiments

Long-term grazing experiments were conducted for all 39 grazers following [24] and 31 were used to quantify grazing rates based on rates of disappearance of *Prochlorococcus* cells, which persist but do not readily grow in K-N medium. Rates were not calculated for eight isolates because they were sampled at a lower frequency. Briefly, flagellate grazers reinoculated from exponentially growing cultures were mixed with live, unstained prey in fresh K-N medium at final concentrations of 10^2^–10^3^ flagellates mL^-1^ and 1.5–2×10^6^ *Prochlorococcus* mL^-1^ and incubated for 3–8 days, depending on how fast prey were ingested. Cell concentrations of prey and grazers were measured every 12–24 hours by flow cytometry on glutaraldehyde-fixed samples (Attune NxT; Thermo Fisher Scientific, Waltham, Massachusetts, USA, and CytoFLEX; Beckman Coulter Life Sciences, Indianapolis, IN, USA). Populations were distinguished based on light scatter and pigment autofluorescence and occasionally confirmed with DNA fluorescence (stained post-sampling with DNA stain SYBR Green I). Ambient bacteria concentrations, monitored using DNA stains, were ≤ 10% of the added *Prochlorococcus* at the start and during most periods of all grazing experiments. Eleven isolates representing all genera (or approximately genus-level clades) were examined in more detail by replicating grazing experiments three times (marked in bold in Supplementary Table S1), while twenty additional isolates were replicated once to better survey within-genus variation.

### Grazing rates and biovolume conversion efficiency

We calculated ingestion rates, *I* (prey grazer^-1^ h^-1^) for each grazer as:

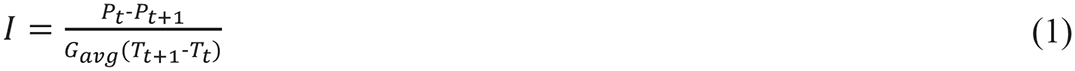

where *P_t_* and *P_t_*_+1_are the prey abundance at sampling interval t and t+1 (cells mL^-1^), *G_avg_* is the arithmetic mean grazer abundance (cells mL^-1^) over the time interval, and (*T_t_*+1 - *T_t_*) is the time (h) between two sampling intervals. Clearance rates (*C*, nL grazer^-1^ h^-1^) were calculated by dividing the ingestion rate by average prey concentration over the same interval, and specific clearance rate (body volume grazer^-1^ h^-1^) was calculated by dividing the clearance rate by cellular biovolume (µm^3^) of each grazer. Final clearance rates reported in the Results were averaged over all sampling intervals (4–8 intervals, 12–24 h for each interval) prior to grazers reaching stationary density.

Six grazers representing three classes (three dictyochophytes, two haptophytes, and one chrysophyte) were further investigated to determine functional grazing responses using a wide range of initial prey densities (10^5^–10^7^ cells mL^-1^). Functional responses were modeled using the Holling type II curve, 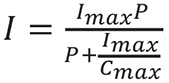, where *I* is the ingestion rate over a sampling interval (eqn. 1),

*I*_max_ is the maximum ingestion rate, *C*_max_ is the maximum clearance rate and *P* is the arithmetic mean prey density between two sampling points. This curve was fit to ingestion rate data using maximum likelihood with R package bbmle [38].

For 31 isolates we calculated the amount of grazer biovolume created per prey biovolume consumed, using data from the same grazing experiments used to calculate grazing rates. It was calculated based on the following formula:

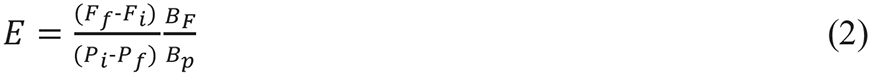

where *F*_f_ and *F*_i_ are the final and initial flagellate concentrations, *P_i_* and *P_f_* are initial and final prey concentrations in each culture, and *B_F_* and *B*_P_ are the cellular biovolume of prey and grazer. We refer to the quantity *E* as the *biovolume conversion efficiency*, and we use it as an indicator of physiological differences among diverse mixotrophs. Note that biovolume conversion efficiency can be greater than 1, if prey have greater nutrient:biovolume than the grazer.

### Quantitative PCR

Real-time, quantitative PCR (qPCR) was performed to quantify the 18S rDNA gene abundances of representative mixotroph groups discriminated at approximately the genus level, including *Florenciella*, *Rhizochromulina* and another undescribed clade within the class Dictyochophyceae; *Chrysochromulina* and another undescribed clade within the class Haptophyceae; Chrysophyceae clade H in the class Chrysophyceae; and certain species of *Triparma* in the class Bolidophyceae (including *T. eleuthera* and *T. Mediterranea* but not *T. Pacifica*). Partial environmental 18S rDNA sequences (700–800 bp) of putative grazers identified in [23] were included for primer design to maximize the potential of targeting various mixotrophic species from each group (Supplementary Fig. S1). Specific primers were designed targeting a short region of 18S rDNA genes for each group ranging between 95–176 bp (Supplement Table S2) and were synthesized by Integrated DNA Technologies (IDT Inc., Coralville, IA, USA).

In situ gene abundances were quantified with water samples collected from Sta. ALOHA by Niskin bottles at 5, 25, 45, 75, 100, 125, 150, and 175m, during HOT cruises #259 (Jan), #262 (Apr), #264 (Jul), and #266 (Oct) of 2014. Seawater (ca. 2 L) was filtered through 0.02 µm pore-sized aluminum oxide filters (Whatman Anotop, Sigma Aldrich, Saint Louis, MO, USA) and stored at −80°C. Genomic DNA of both grazer cultures and environmental samples were extracted (MasterPure Complete DNA and RNA Purification Kit; Epicentre) according to [39]. Four replicated PCR reactions (10 µL) were carried out for each sample except for *Triparma* (duplicates) and consisted of 5 µl of 2× PowerTrack SYBR Green Master Mix (Thermo Fisher Scientific, USA), 10 ng environmental DNA, 500 nM of each primer, and nuclease-free water. Reactions were run on an Eppendorf Mastercycler epgradient S realplex^2^ real-time PCR instrument. Each run contained fresh serial dilutions (1–6 log gene copies µL^-1^) of target-specific, 750-bp synthetic standards (gBlocks, IDT) prepared in triplicate. The cycling program included an initial denaturation step of 95 °C for 2 min, followed by 40 cycles of 95 °C for 5 s and 55 °C for 30 s. Specificity of amplification was checked with a melting curve run immediately after the PCR program and gel electrophoresis. Amplification efficiencies ranged from 97% to 104% for all the primers.

To allow conversion from gene copies to cell numbers, the 18S rDNA gene copy number per cell was determined for the seven targeted clades. Known amounts of cultured cells (10^6^–10^7^ cells) were pelleted at 4,000 RCF for 15 min at 4 °C, DNA was extracted, and the gene copy numbers were determined with the group-specific primers. Cell abundances in both original cultures and supernatants were monitored by flow cytometry to account for losses. Various isolates from each group were selected for this purpose and a mean cellular 18S rDNA gene copy number was calculated for each genus. To estimate contributions of each isolate to the local community of small phytoplankton at Station ALOHA, calculated in situ cell abundances were compared to counts of photosynthetic picoeukaryotes from the Hawaiʻi Ocean Time-series (https://hahana.soest.hawaii.edu/hot/hot-dogs/).

### Global distribution revealed through Tara Oceans 18S rDNA metabarcodes

To estimate the relative abundance of the OTUs closely related to our diverse isolates on a broader geographic scale, we searched the 18S rDNA-V9 sequence data from the 0.8–5 µm fraction of surface water sampled at 40 stations by the *Tara* Oceans project (http://taraoceans.sb-roscoff.fr/EukDiv/). Reads for a ‘*Tara* lineages’ with highest similarity (E-value < 10^-15^) to each of our targeted clades (Supplementary Table S1) were expressed as a fraction of total reads excluding dinoflagellates but included all other *Tara* Oceans phytoplankton ‘taxogroups’: Bacillariophyta, Bolidophyceae, Chlorarachnea, Chlorophyceae, Chrysophyceae/Synurophyceae, Cryptophyta, Dictyochophyceae, Euglenida, Glaucocystophyta, Haptophyta, Mamiellophyceae, Other Archaeplastida, Other Chlorophyta, Pelagophyceae, Phaeophyceae, Pinguiophyceae, Prasino-Clade-7, Pyramimonadales, Raphidophyceae, Rhodophyta and Trebouxiophyceae. Dinoflagellates were excluded because of the difficulty in assigning phototrophic vs. heterotrophic status to all taxa, and because nearly all dinoflagellate reads were from a single, poorly annotated OTU that was also highly abundant in larger size fractions.

## Results

### Phylogenetic diversity

Diverse *Prochlorococcus*-consuming mixotrophic flagellates ranging in size from 2–5 µm were isolated from oligotrophic, open-ocean waters of the NPSG (Table 1). The isolates include species from nine genera (or approximately genus-level clades, referred to hereafter as genera for brevity) and six classes (Fig. 1a; Supplementary Table S1): one *Pelagodinium* isolate in class Dinophyceae; one isolate in an undescribed clade within class Chlorarachniophyceae (hereafter referred to as ChloraX); two *Chrysochromulina* isolates and one environmental HAP-2 clade isolate (hereafter Hap2) [40] in class Haptophyceae; three environmental clade H isolates in class Chrysophyceae (hereafter ChrysoH) [41]; two *Triparma* isolates in class Bolidophyceae; and twenty-nine isolates in class Dictyochophyceae: nineteen *Florenciella* isolates, four *Rhizochromulina* isolates, and six isolates in an undescribed clade (hereafter DictyX). The *Florenciella* isolates cluster with two cultivated strains (*Florenciella parvula*, GenBank Acc. AY254857; Florenciellales sp. NIES 1871, GenBank Acc. AB518483) and one environmental sequence (GenBank Acc. AY129059). The three chrysophyte isolates are closely related to an environmental Chrysophyceae clade H sequence (GenBank Acc. EF172998). The closest relatives of the two *Triparma* isolates are *Triparma eleuthera* and *Triparma mediterranea*, and the two *Chrysochromulina* isolates are most closely related to an environmental *Chrysochromulina* (GenBank Acc. AF107083) and *Chrysochromulina* sp. RCC400 (GenBank Acc. KT861300), respectively. All isolates showed evidence of phagocytosis (e.g., Fig. 1b) and maintained permanent chloroplasts.

**Fig. 1.**
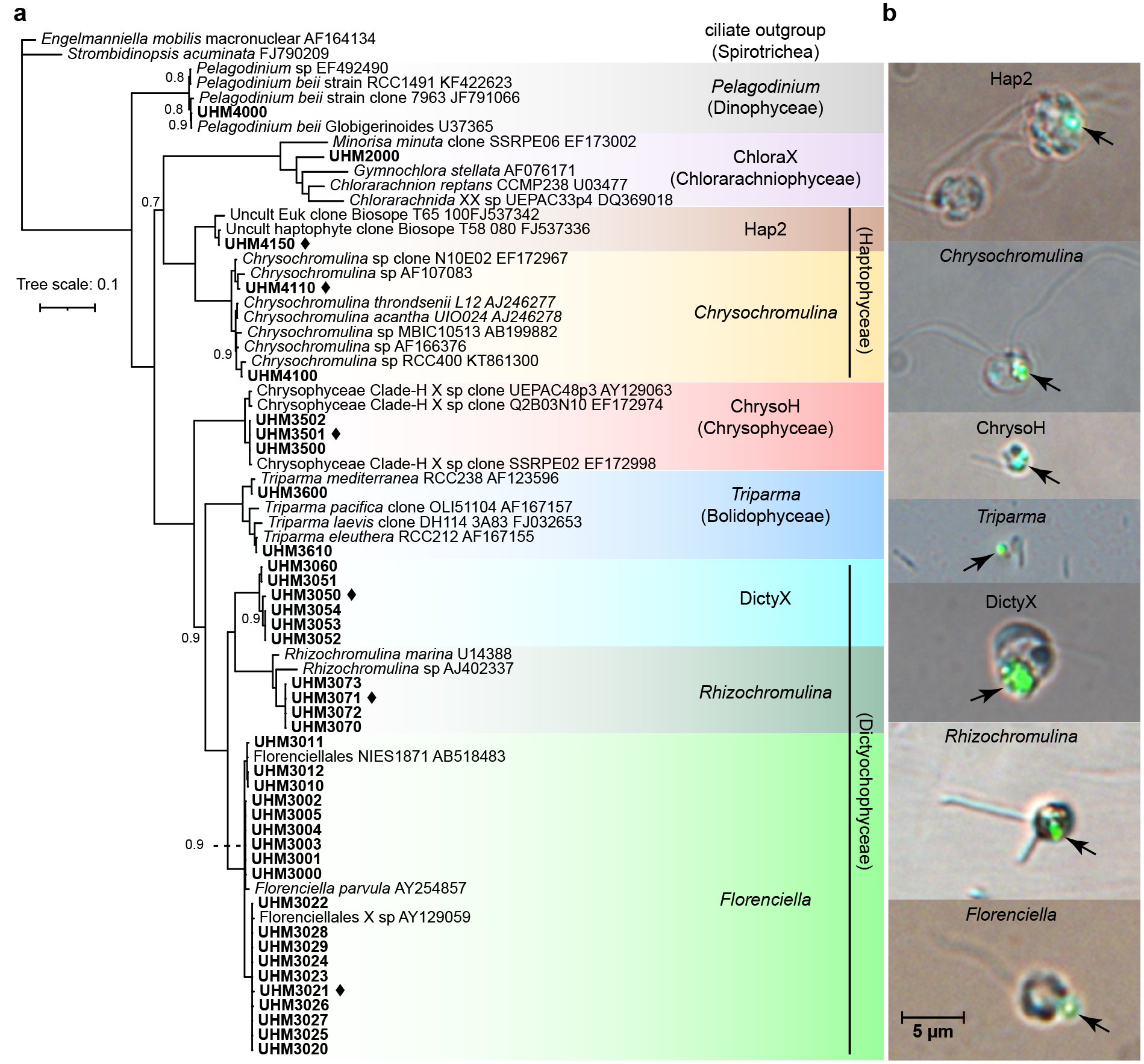
Phylogenetic diversity among the isolates and visual evidence of phagotrophy. (**a**) A near-full length 18S rDNA-based Bayesian majority consensus tree, showing the phylogenetic positions of 39 mixotrophic grazers isolated from the NPSG (in bold). Genera are shaded with different colors and labeled with genus and class (in parentheses) names, and the six grazers investigated for grazing functional responses are marked with black diamonds. Most branches have complete support (posterior probability = 1.0) and only those with lower support (<1.0) are labeled with support values. The scale bar indicates 10% divergence. (**b**) Digital micrographs of grazers consuming 0.5-µm fluorescent beads were captured using transmitted light and epifluorescence microscopy. The scale bar is 5 µm.

### Grazing capability and growth efficiency

All 39 isolates were confirmed to consume *Prochlorococcus* and grew when *Prochlorococcus* was the sole added prey and primary source of nitrogen. Within a five-day time course, consumption of 1–2×10^6^ *Prochlorococcus* mL^-1^ supported grazer growth of 10^3^–10^4^ cells mL^-1^ for the different isolates (Supplementary Fig. S2). Grazing rates varied across phylogenetic groups, with >10-fold and ∼6-fold variation in both clearance rate (volume cleared per cell) and specific clearance rate (volume cleared per body volume) across species and genera, respectively (Fig. 2a–b). Isolate UHM3050 of clade DictyX possessed the highest raw clearance rate of ∼10 nL grazer^-1^ h^-1^ and strain UHM3501 of clade ChrysoH displayed the highest specific clearance rate of ∼5×10^5^ body volumes grazer^-1^ h^-1^. On the genus level, ChrysoH also had the highest specific clearance rate (3.6×10^5^ body volumes grazer^-1^ h^-1^), followed by three clades of *Triparma*, *Rhizochromulina*, and DictyX (1.6–1.9×10^5^ body volumes grazer^-1^ h^-1^) and *Chrysochromulina* (1.2×10^5^ body volumes grazer^-1^ h^-1^). *Florenciella* are among the lowest in terms of both raw and specific clearance rates (1.2 nL grazer^-1^ h^-1^, 0.5×10^5^ body volumes grazer^-1^ h^-1^). The remaining flagellates showed clearance rates in between these extremes, with significant variation in rates explained by genus (ANOVA on log-transformed clearance rates: F_9,20_ = 8.8, p < 10^-4^, R^2^ = 0.80; ANOVA on log-transformed specific clearance rates: F_9,20_ = 7.6, p < 10^-4^, R^2^ = 0.77).

**Fig. 2.**
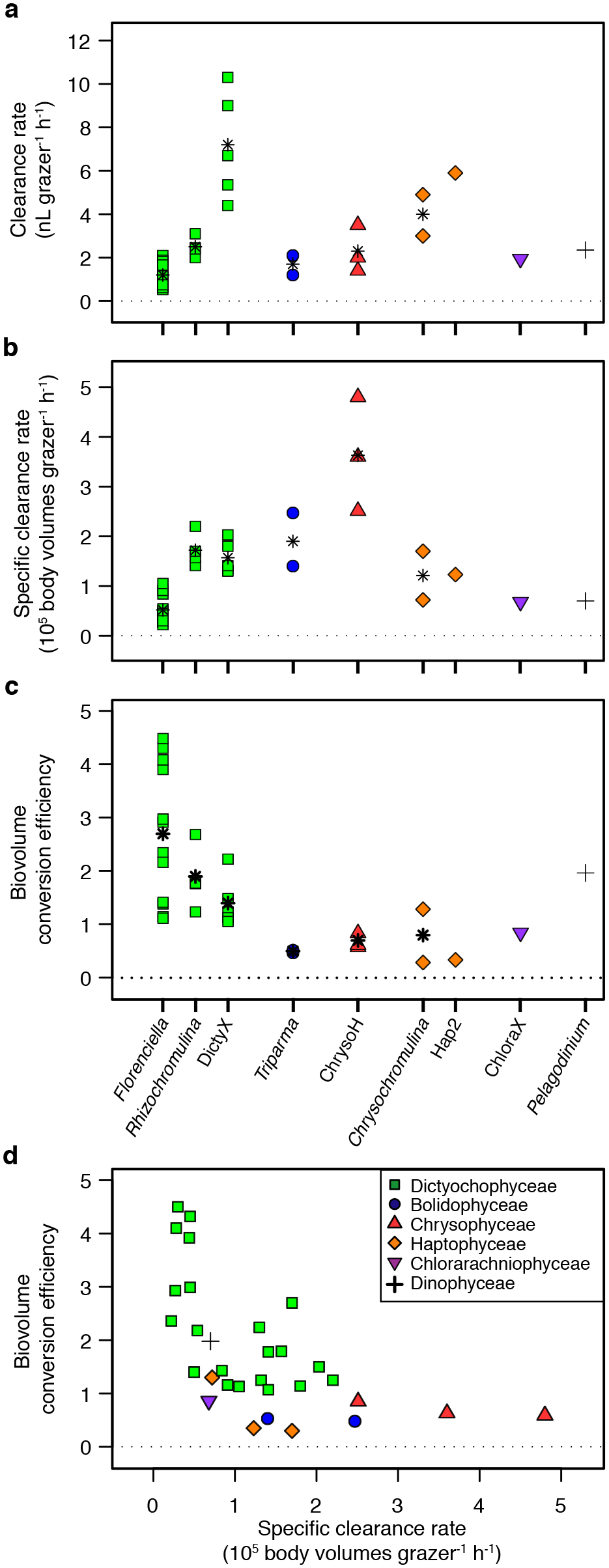
Comparison of grazing rates and growth efficiencies among NPSG mixotrophs. Panels show (**a**) clearance rates or (**b**) specific clearance rates among isolates in different genera and biovolume conversion efficiencies for isolates (**c**) grouped by genus or (**d**) as a function of specific clearance rates. Individual classes were denoted with different colors and symbols in all four panels (legend in panel **d**), and genera that have multiple strains included an arithmetic mean shown in black asterisk. Note that for some isolates (marked in bold in Supplementary Table S1) replicated grazing experiments were conducted and the means are presented in this figure.

Biovolume conversion efficiency also varied among genera, with the highest mean value observed for *Florenciella* (2.7), followed by the other two dictyochophyte clades, *Rhizochromulina* (1.9) and DictyX (1.4). The lowest efficiencies were exhibited by the two *Triparma* (0.5), three ChrysoH (0.7) and three haptophyte isolates (0.8) (Fig. 2c). Strains with higher specific clearance rates tended to have lower growth efficiencies (r = −0.64, p < 10^-3^; Fig. 2d). There was substantial variation in these traits even within the Dictyochophyceae, with the highest conversion efficiencies seen in *Florenciella* strains that possessed the lowest clearance rates (e.g., UHM3029 and UHM3023), and some of the lowest conversion efficiencies found in strains with higher clearance rates (e.g., *Rhizochromulina* UHM3071 and DictyX UHM3052).

### Functional responses

Functional responses were estimated for six strains representing different classes and genera (Fig. 3; strains denoted in Table 1 and Fig. 1). Maximum ingestion rates (*I*_max_) ranged from 2 cells grazer^-1^ h^-1^ (ChrysoH) to 92 cells grazer^-1^ h^-1^ (Hap2), with varying prey saturation concentrations between 10^5^ cells mL^-1^ for ChrysoH, 10^6^ cells mL^-1^ for *Rhizochromulina* and 10^7^ cells mL^-1^ for the remaining strains (Fig. 3). Specific *I*_max_ (volume ingested per body volume) ranged from 0.04 body volumes grazer^-1^ h^-1^ (ChrysoH) to 0.43 body volumes grazer^-1^ h^-1^ (*Chrysochromulina*), respectively and specific *C*_max_ ranged from 0.9×10^5^ body volumes grazer^-1^ h^-1^ (*Florenciella*) to 1.9×10^6^ body volumes grazer^-1^ h^-1^ (ChrysoH) (Supplementary Table S3). Substantially higher growth of grazers and faster removal of prey were supported by higher *Prochlorococcus* densities (e.g., 5×10^6^–5×10^7^ cells mL^-1^), compared to treatments with lower *Prochlorococcus* densities (< 1×10^6^ cells mL^-1^) (Supplementary Fig. S3).

**Fig. 3.**
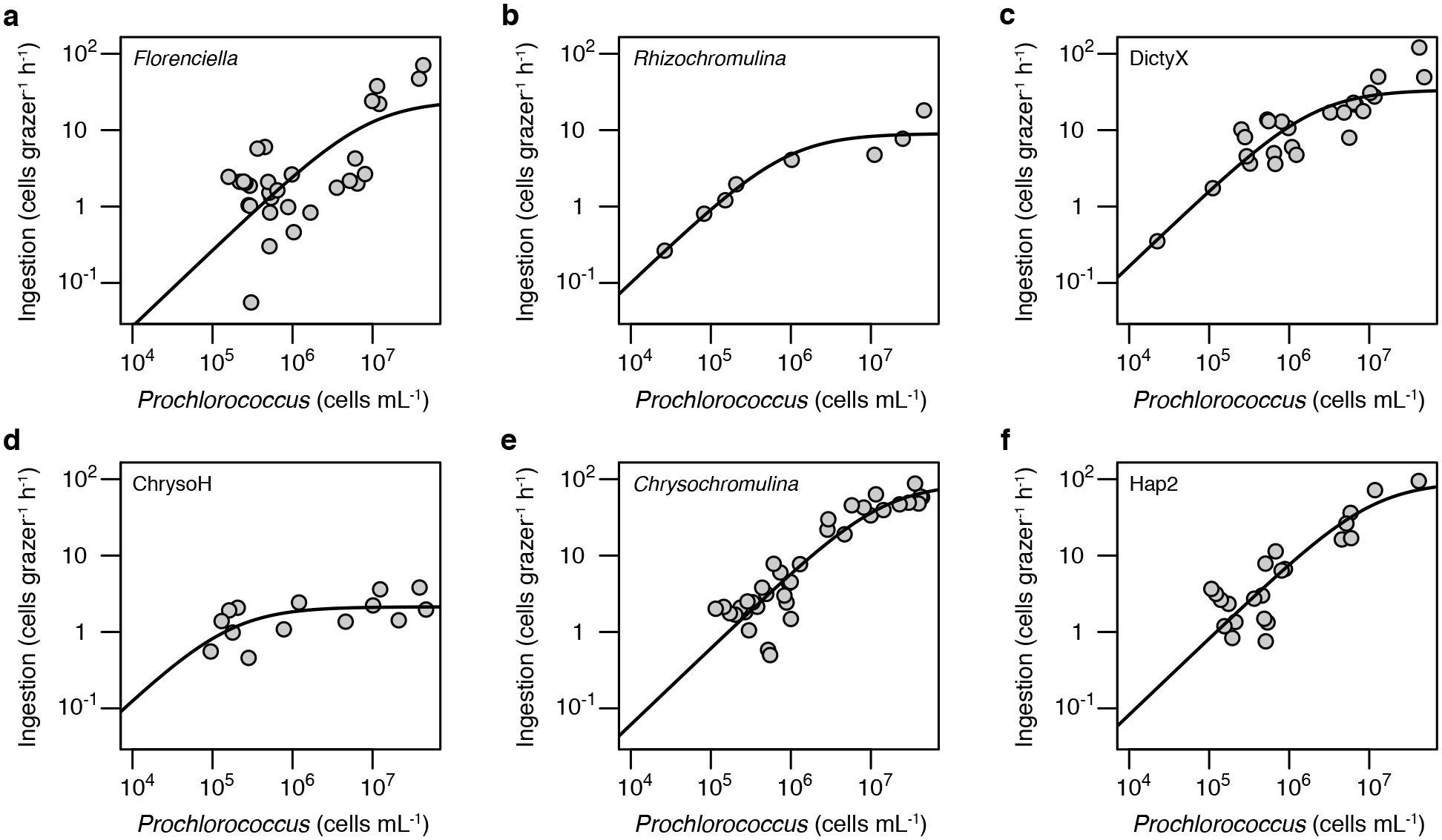
Functional responses of six representative grazers from different genera. Ingestion rates of *Prochlorococcus* are plotted as a function of prey concentration. Holling type II curves were fitted to the data to estimate *C*_max_ (maximum clearance rate, the initial slope of the curve) and *I*_max_ (maximum ingestion rate, the asymptote of the curve). Grazer and prey trajectories from experiments used to estimate functional responses are shown in Supplementary Fig. S3.

The body volume-specific *C*_max_ and *I*_max_ of our isolates cover nearly the whole range of values previously reported in the literature for nano-sized heterotrophic and mixotrophic flagellates, which are from <10^4^ to >10^6^ body volumes grazer^-1^ h^-1^, and <0.1 to >1 body volumes grazer^-1^ h^-1^, respectively (Fig. 4; Supplementary Table S3). It is noteworthy that the ChrysoH strain studied here, which is the smallest of our isolates (2–3 µm), demonstrated a higher specific *C*_max_ than any protistan grazer studied to date. For the one isolate previously studied and re-examined here (*Florenciella* UHM 3021), grazing rates were somewhat faster than previously measured, but both prior and current *C*_max_ estimates for this strain are substantially lower than the other five isolates analyzed (Supplementary Table S3), consistent with the general pattern of low clearance rates for this genus (Fig. 2).

**Fig. 4.**
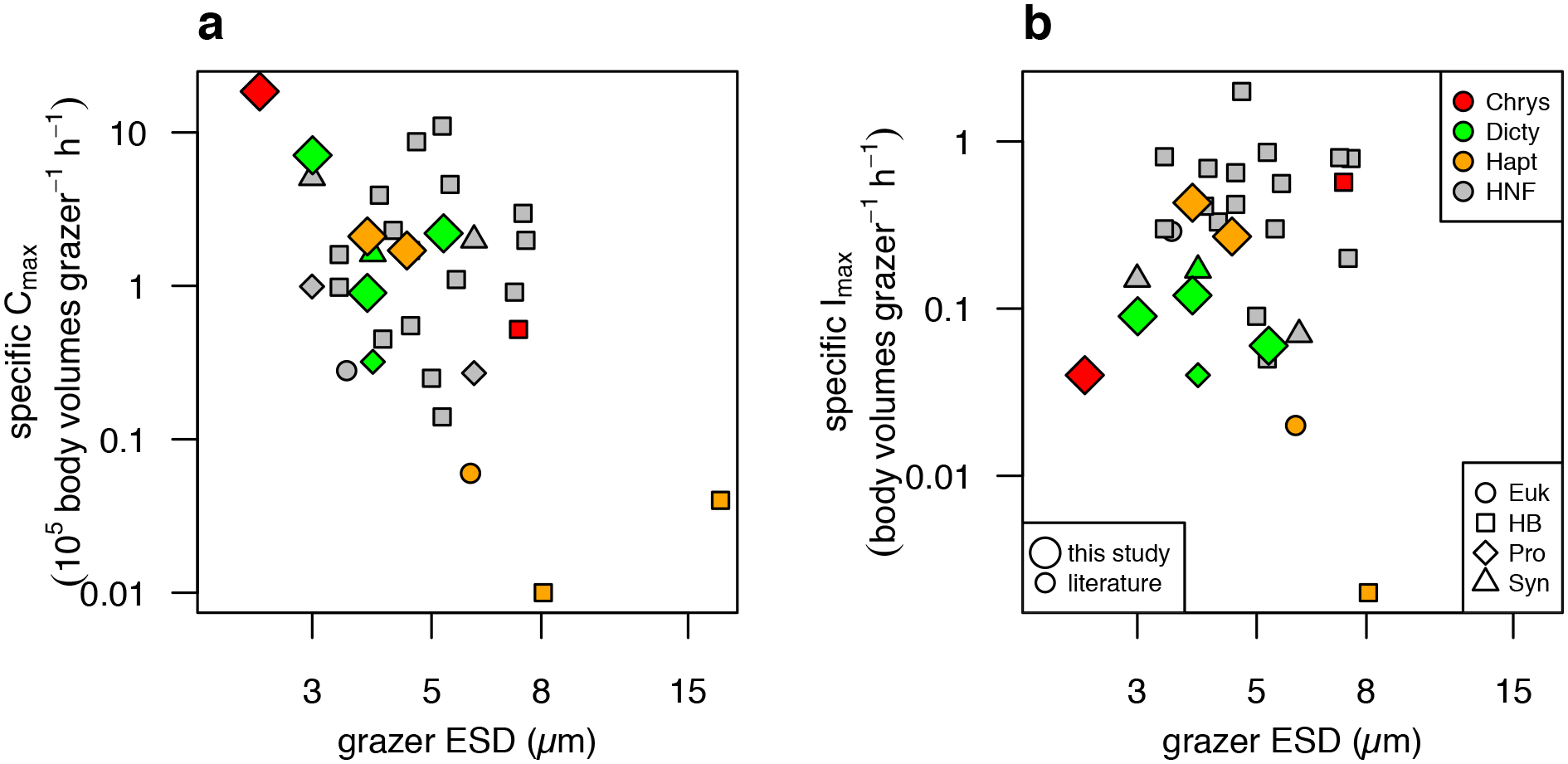
Grazing rate parameters as a function of size (equivalent spherical diameter, or ESD) for heterotrophic (HNF) and mixotrophic nanoflagellates that have been studied in culture. (**a**) Specific *C*_max_ (maximum volume cleared per body volume) and (**b**) specific *I*_max_ (maximum cell volume ingested per body volume), for flagellates grazing on *Prochlorococcus* (Pro), *Synechococcus* (Syn), heterotrophic bacteria (HB), or eukaryotic prey (Euk). In both panels (legend in panel b), grey-filled symbols indicate HNF grazers, and other color-filled symbols indicate mixotrophic grazers (both from this study and literature), including chrysophytes (Chrys), dictyochophytes (Dicty) and haptophytes (Hapt). Symbol shapes indicate the type of prey (source data in Supplementary Table S3). Large symbols indicate data from this study.

### Abundance in the field

Estimates of 18S rDNA gene copy numbers indicate that all isolates tend to have low copy numbers per cell, with approximately one copy cell^-1^ for ChrysoH and *Triparma*, one to three copies cell^-1^ for *Florenciella* and *Rhizochromulina*, two copies cell^-1^ for *Chrysochromulina*, three copies cell^-1^ for Hap2, and five copies cell^-1^ for DictyX (Fig. 5a). 18S rDNA gene copy number correlated with cell size among these isolates either at the strain level (r = 0.78, p < 0.001), or when grouped by genus (r = 0.90, P < 0.01) (Fig. 5b).

**Fig. 5.**
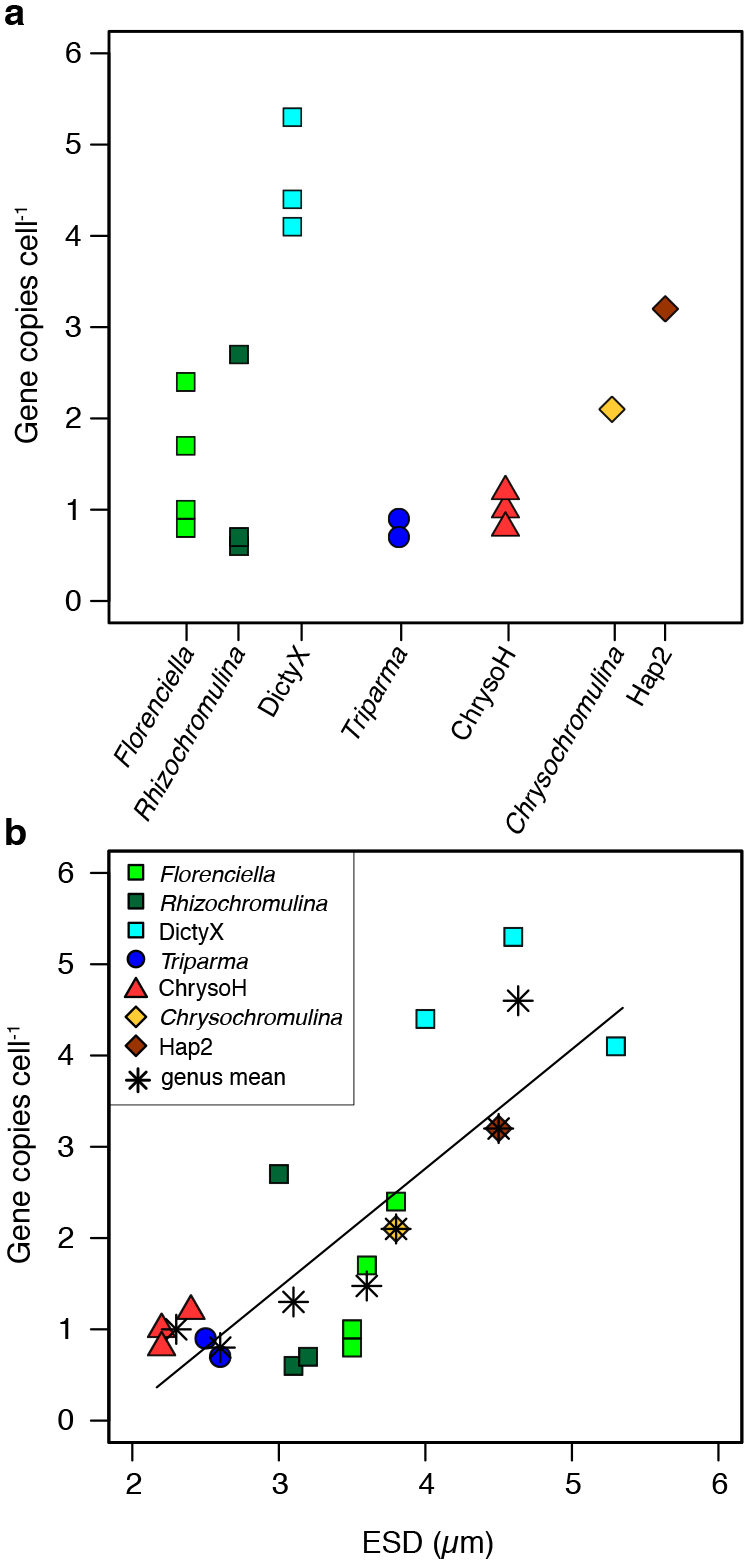
18S rDNA gene copy numbers per cell, shown for (**a**) each of seven genera or (**b**) as a function of equivalent spherical diameter (ESD) of the cells. Each colored symbol in both panels denotes the mean of a single isolate (averaged over triplicate measurements) and asterisk symbols in panel (**b**) are mean values for each genus (averaged over isolates). Least-squares linear regression line is plotted for isolate means (R^2^ = 0.61).

Average abundance of 18S rDNA gene copies in the euphotic zone at Sta. ALOHA varied from 1×10^2^ to 5.3×10^5^ copies L^-1^ across groups, and estimated cell concentrations varied from 1×10^2^ to 2.5×10^5^ cells L^-1^ (Fig. 6). All groups were more abundant at upper euphotic depths (5– 100 m), with abundance typically peaking at 45 m, except for *Florenciella* (75 m) and *Chrysochromulina* (100 m). *Chrysochromulina* was most abundant at every depth, peaking at 5.3×10^5^ copies L^-1^ (2.5×10^5^ cells L^-1^) at 100 m. The Hap2 clade had the second highest gene abundance (1.2×10^5^ copies L^-1^ at 45 m), but estimated cell abundances of Hap2, *Florenciella* and ChrysoH clades were all very similar with maxima of 3.3–3.6×10^4^ cells L^-1^ at 45–75 m. Lower abundances were seen for *Triparma*, *Rhizochromulina*, and DictyX clades, which had maxima ranging between 1.1–1.5×10^4^ copies L^-1^ (3×10^3^ –1.3×10^4^ cells L^-1^) at 45 m.

**Fig. 6.**
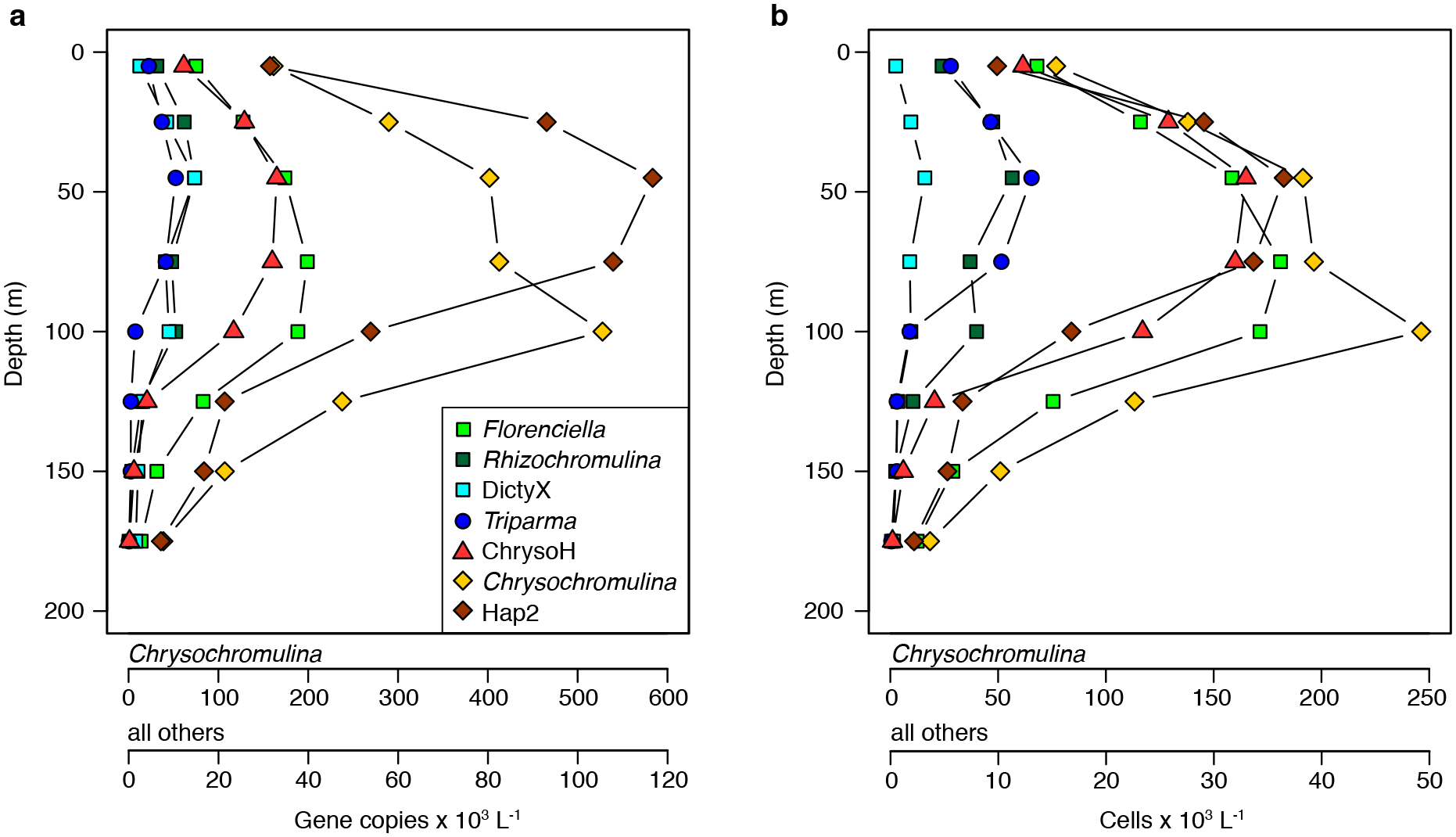
Average in situ abundance as a function of depth of (**a**) 18S rDNA gene copies and (**b**) the corresponding inferred cell abundances. Results are the averages from four depth profiles (one collected in each of four seasons during 2014). Complete source data for each season shown in Supplementary Fig. S4.

Estimated contributions of each clade to the total photosynthetic picoeukaryotes in upper euphotic depths (0–100 m) were 0.1–1% (DictyX, *Rhizochromulina* and *Triparma*), 1–4% (ChrysoH, *Florenciella* and Hap2) and 9–23% (*Chrysochromulina*) (Supplementary Fig. S4a). In total these targeted clades accounted for a relative abundance between 14%–31% in upper (0– 100 m) and 7–14% in lower euphotic depths (125–175 m), respectively. Group abundances varied over time (Supplementary Fig. S4b–h), with DictyX (Supplementary Fig. S4d) and Hap2 (Supplementary Fig. S4g) clades more abundant during October, while *Triparma* (Supplementary Fig. S4e) and ChrysoH (Supplementary Fig. S4f) clades were more abundant during April.

Analysis of *Tara* Oceans OTUs closely related to our isolates indicates that these taxa are widespread across surface ocean samples (Fig. 7 a-c), with median relative abundances ranging from < 0.1% to around 5% (Fig. 7d). Dictyochophyte and haptophyte each constituted 10-25% of the community at over 10 stations, and their most abundant groups of Florenciella parvula and Chrysochromulina_X sp. alone accounted for > 10% at 5 stations (Fig. 7 a-b). In total the focal OTUs accounted for 5–36% (median 15%) of non-dinoflagellate phytoplankton in the 0.8–5 µm size fraction (Fig. 7d).

**Fig. 7.**
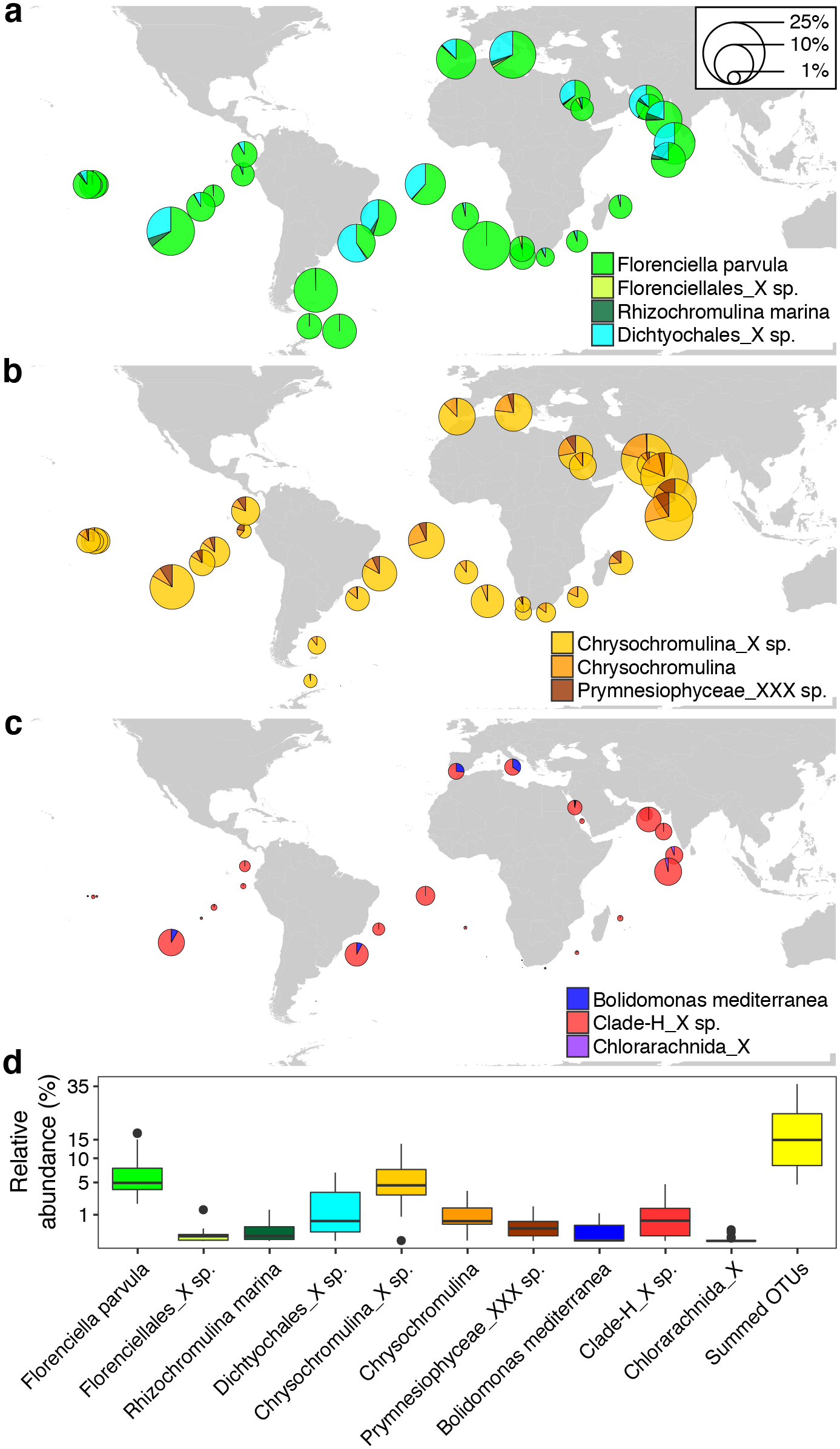
Global patterns of *Tara* Oceans OTUs closely related to our mixotroph isolates. Relative abundances of ten OTUs were grouped into (**a**) Dictyochophyceae, (**b**) Haptophyceae, and (**c**) others, and are represented with pie charts at each station or (**d**) presented as box and whisker plots of the percent contributions for each targeted OTU (n = 40). Outliers determined by the 1.5×IQR rule are shown as filled circles. Pie chart areas are proportional to OTU relative abundances expressed as a percentage of total phytoplankton OTUs (excluding dinoflagellates) in the 0.8-5 µm size fraction. OTUs are labeled with their lowest-level taxonomic annotations from the *Tara* Oceans dataset (full *Tara* lineages, numeric IDs, and corresponding isolates are presented in Supplementary Table S1).

## Discussion

### Phylogenetically diverse and globally common mixotrophs

Marine protists are incredibly diverse and much of this diversity remains uncultured, even for the relatively well-studied phytoplankton, which makes it challenging to interpret the functional significance of environmental sequence data. The flagellates isolated and characterized in this study appear to be the only mixotrophs for which consumption of *Prochlorococcus* has been confirmed and quantified in culture-based lab experiments.

Quantitative data on the grazers of *Prochlorococcus* is important because these cyanobacteria are major primary producers in oligotrophic waters and the most abundant autotroph on a global scale [13][42]. Among the mixotrophic flagellates described here are representatives of four genera (DictyX, ChrysoH, Hap2, ChloraX) from which no cultivated representatives have previously been reported, but which appear frequently in molecular surveys. We have quantified grazing behavior in these isolates, as well as isolates from three described genera (*Rhizochromulina*, *Triparma,* and *Pelagodinium*) that had not previously been documented to consume prey. We have also studied isolates of *Chrysochromulina* from the oligotrophic open ocean, whereas previous isolates were mostly from productive coastal environments and algal bloom events. Our isolates are smaller than most previously studied mixotrophs, perhaps reflecting their origins in the subtropical oligotrophic open ocean (Supplementary Table S4).

Comparison of our isolates to environmental sampling efforts suggests that *Prochlorococcus*-consuming mixotrophs are common components of open ocean ecosystems, with two genera of *Florenciella* and *Chrysochromulina* being ubiquitous in certain regions (Fig. 7a-b). The aggregate relative abundance of OTUs related to our isolates in surface *Tara* Oceans contributed up to 36% and the median abundance is similar to qPCR-based relative abundance of the targeted clades at Sta. ALOHA, i.e., 15% vs. 14% (Supplementary Fig. S4a; Fig. 7). This suggests that our isolates may represent the most abundant populations in these clades, and further efforts to estimate the absolute abundance of these globally-common mixotrophs are necessary.

### A broad spectrum of trophic strategies

The NPSG is one of the most oligotrophic ecosystems in the global ocean, with persistently depleted surface nutrients and relatively stable microbial distributions [44]. The dominant eukaryotic phototrophs are small, and the 2–5 µm size class alone contributes approximately half of eukaryotic phytoplankton biomass and about one-fifth of total phytoplankton biomass [22]. This community includes non-trivial populations of haptophytes, dinoflagellates, dictyochophytes, chrysophytes, prasinophytes, bolidophytes, cryptophytes, and pelagophytes [1][43]. Our results suggest that the high diversity of these organisms may be attributable in part to a broad spectrum of trophic strategies, including autotrophy (e.g., immotile prasinophytes) and a variety of mixotrophic strategies, illustrated by the large variation in grazing abilities among our isolates. Differences among genera in the traits we have measured suggests that the functional capacity of these communities may be somewhat predictable based on taxonomic information alone. We hypothesize that fast grazers such as the ChrysoH clade are relatively heterotrophic mixotrophs, acquiring more energy from prey and less from photosynthesis compared to slower grazers such as *Florenciella*. Although we do not have data on autotrophic capacities, a spectrum of autotrophy vs. heterotrophy is consistent with a tradeoff between clearance rate and biovolume conversion efficiency (Fig. 2d), because relatively autotrophic mixotrophs may incorporate more prey biomass into grazer biomass than relatively heterotrophic mixotrophs that respire more prey biomass for energy [45][46]. Therefore, retention vs. remineralization of prey biomass may vary substantially across mixotrophs that co-occur in the same ecosystem, and the consequences for aggregate nutrient cycling and trophic transfer efficiency will depend on which physiologies are most abundant.

Maximum ingestion rates of the NPSG mixotrophs are intermediate or high compared to previously studied mixotrophs including haptophytes and chrysophytes, but are generally lower than most heterotrophs (Fig. 4). These results are consistent with the expectation that mixotrophs are constrained in their phagotrophic ability by tradeoffs with functions such as photosynthesis and nutrient uptake. At the same time, maximum clearance rates of the NPSG mixotrophs are intermediate-to-high compared to heterotrophs, suggesting that the two traits (*I*_max_ and *C*_max_) may be constrained by different mechanisms. *C*_max_ quantifies grazing performance when grazing is encounter-limited, and it may reflect swimming speed or feeding currents that determine encounter rate, or the efficacy of prey capture per encounter [47]. In contrast, *I*_max_ may reflect the rate of vacuole formation or other processes limiting the rate of digestion. Comparison of such traits among diverse mixotrophs, as well as phototrophic traits, would illuminate the physical basis of differences in grazing ability.

### In situ abundances and potential grazing impacts

At Station ALOHA, a typical abundance of small pigmented flagellates is ∼1000–2000 cells mL^-1^ [22][48], and many of these photosynthetic picoeukaryotes may be mixotrophic based on RNA stable isotope labeling experiments [23]. However, counts of individual taxa of small protists are scarce [49]. Our qPCR estimates show that the haptophyte genus *Chrysochromulina* was the most abundant of the groups we targeted, and the targeted groups together accounted for up to ∼30% of the flow cytometric counts of total photosynthetic picoeukaryotes. A recent study at this location using fluorescently labeled bacterial prey estimated mixotrophs at 5–25 m to be ∼100–200 cells mL^-1^ [17]. This is comparable to the sum of our targeted mixotroph taxa, which suggests the targeted groups could account for most of the mixotrophs. However, use of labeled prey may underestimate the total grazer population and our assays did not target all potential mixotrophs, so the contribution of mixotrophs to total photosynthetic eukaryotes is likely > 30%. Based on a previous metabarcoding study at this location [43], the remaining small, pigmented eukaryotes likely include presumed autotrophs (immotile prasinophytes, coccolithophores, *Pelagomonas*), but also other presumed mixotrophs (dinoflagellates, cryptophytes, additional clades within the classes that our mixotrophs belong to) that we did not target (Supplementary Fig. S1).

When combining estimated clade abundances with the maximal clearance rates of our isolates (2.7–16.6 nL flagellate^-1^ h^-1^), we predict that the targeted mixotrophs could consume up to 15% of *Prochlorococcus* produced daily in upper euphotic depths (0–100 m) at Sta. ALOHA, using *Prochlorococcus* production estimates at this site from [50]. If we assume that all pigmented picoeukaryotes in this system are mixotrophs, and that they graze with a clearance rate of 6.6 nL flagellate^-1^ h^-1^ (weighted average of different mixotroph groups), then grazing mortality by mixotrophs would account for 26–50% of daily *Prochlorococcus* production in upper euphotic depths. These results suggest bacterial removal through mixotrophic grazing is an important process for mortality. However, the true grazing contribution from mixotrophs will depend on the relative abundance of autotrophs vs. mixotrophs, a number that remains elusive, as well as mixotroph community structure, as we have found that grazing abilities vary substantially among taxa. Furthermore, to better understand mixotrophy in ocean ecosystems it will be important to test how the prevalence of different trophic strategies is related to environmental conditions, and whether this leads to predictable gradients in how mixotrophs affect productivity and nutrient cycling.

## Acknowledgments

This work was supported by National Science Foundation awards OCE 15-59356 and OIA 17- 36030, and a Simons Foundation Investigator Award in Marine Microbial Ecology and Evolution (to K.F.E.). We thank the personnel in the Hawai’i Ocean Time-series program (NSF award 12-60164) for assistance with water sampling. We thank Dr. Sallie W. Chisholm (Massachusetts Institute of Technology) for providing the *Prochlorococcus* strains. We are also grateful to Tina M. Weatherby at the Biological Electron Microscope Facility of the University of Hawaiʻi at Manoa (UHM), as well as Dr. Karen E. Selph at the School of Ocean and Earth Science and Technology of UHM for their assistance with microscopic imaging and flow cytometry measurements.

## Competing interests

All authors declare that they have no competing interests related to this work.

## Supplementary Material

**Supplementary Fig. S1.**
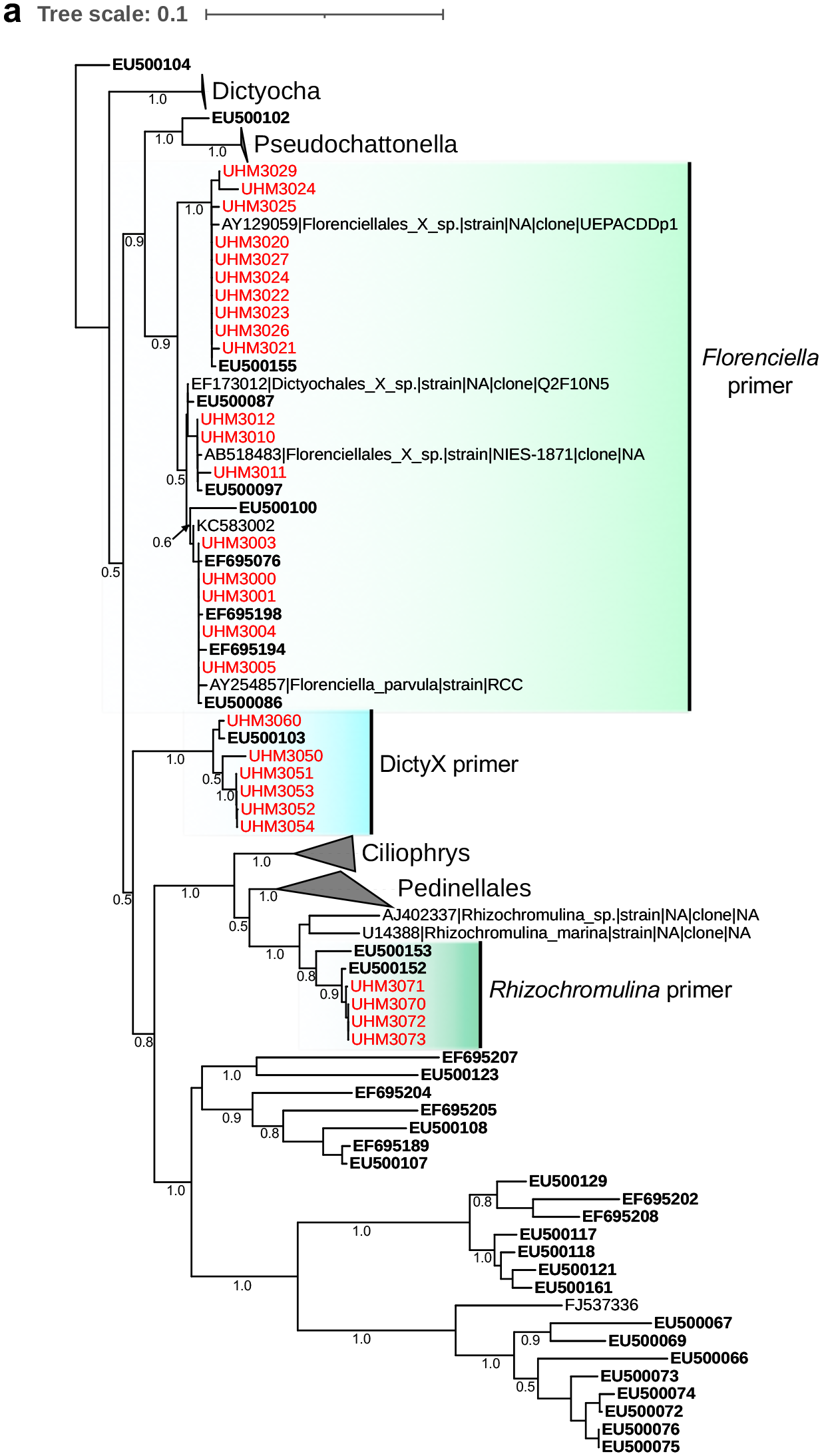
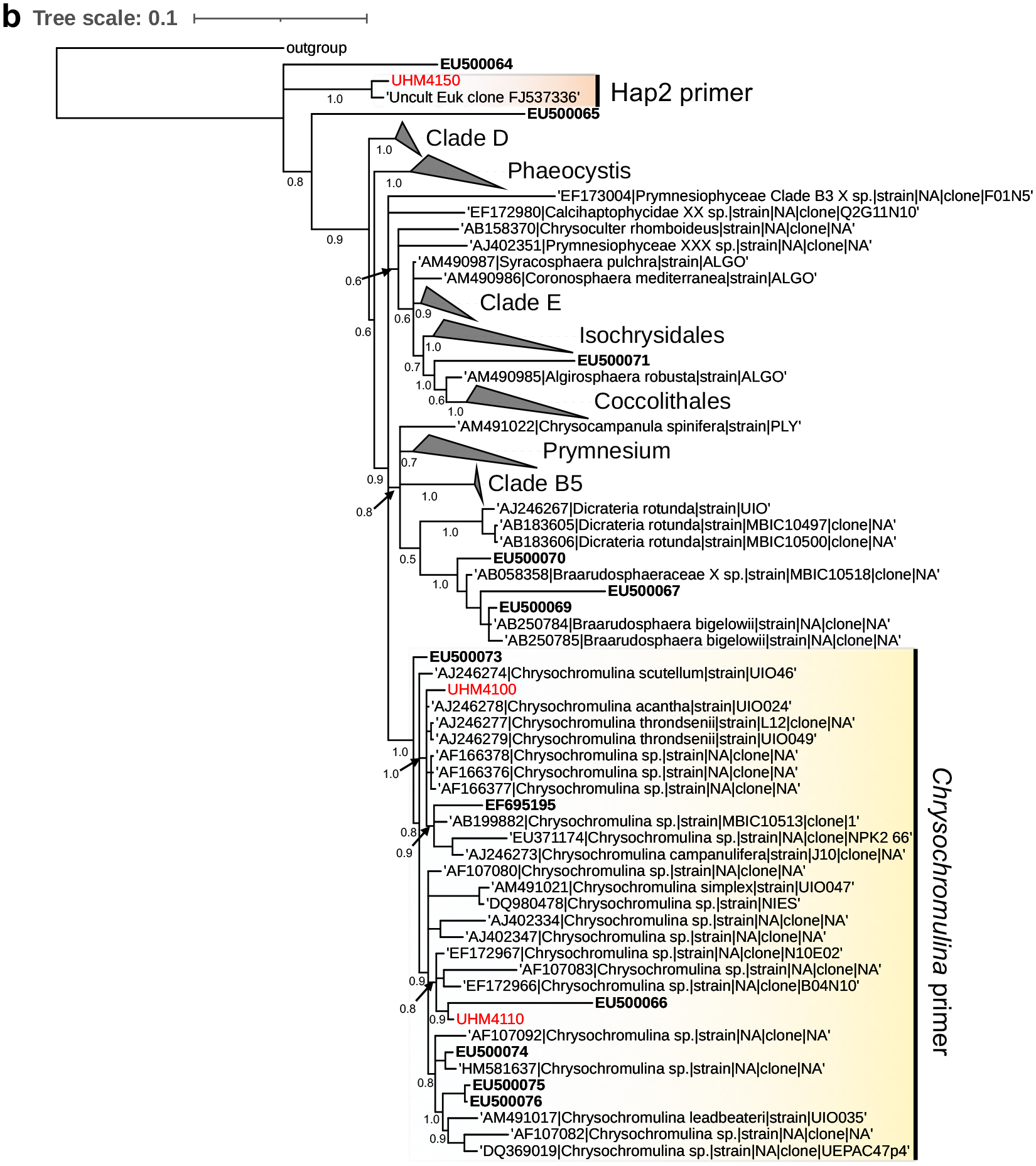
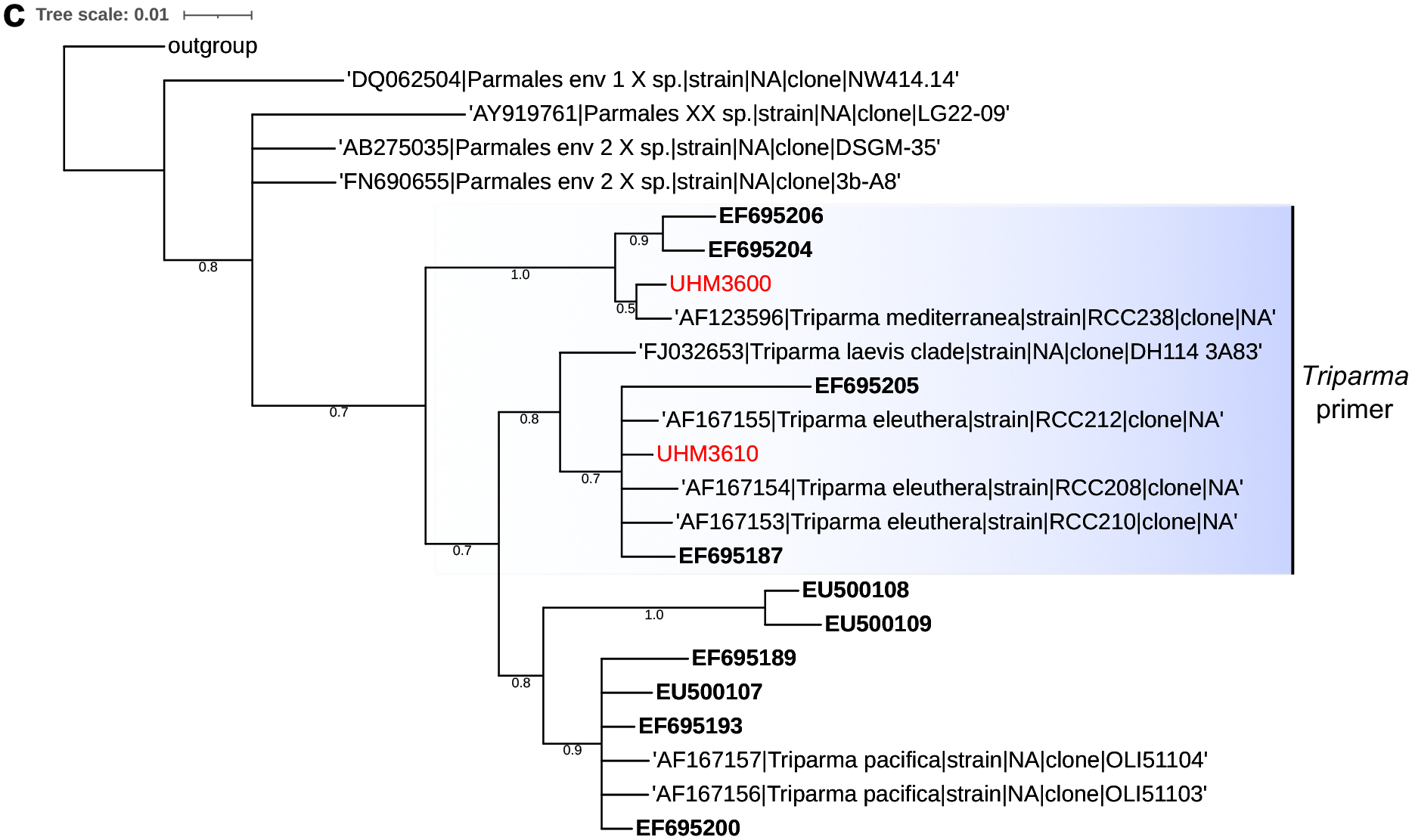
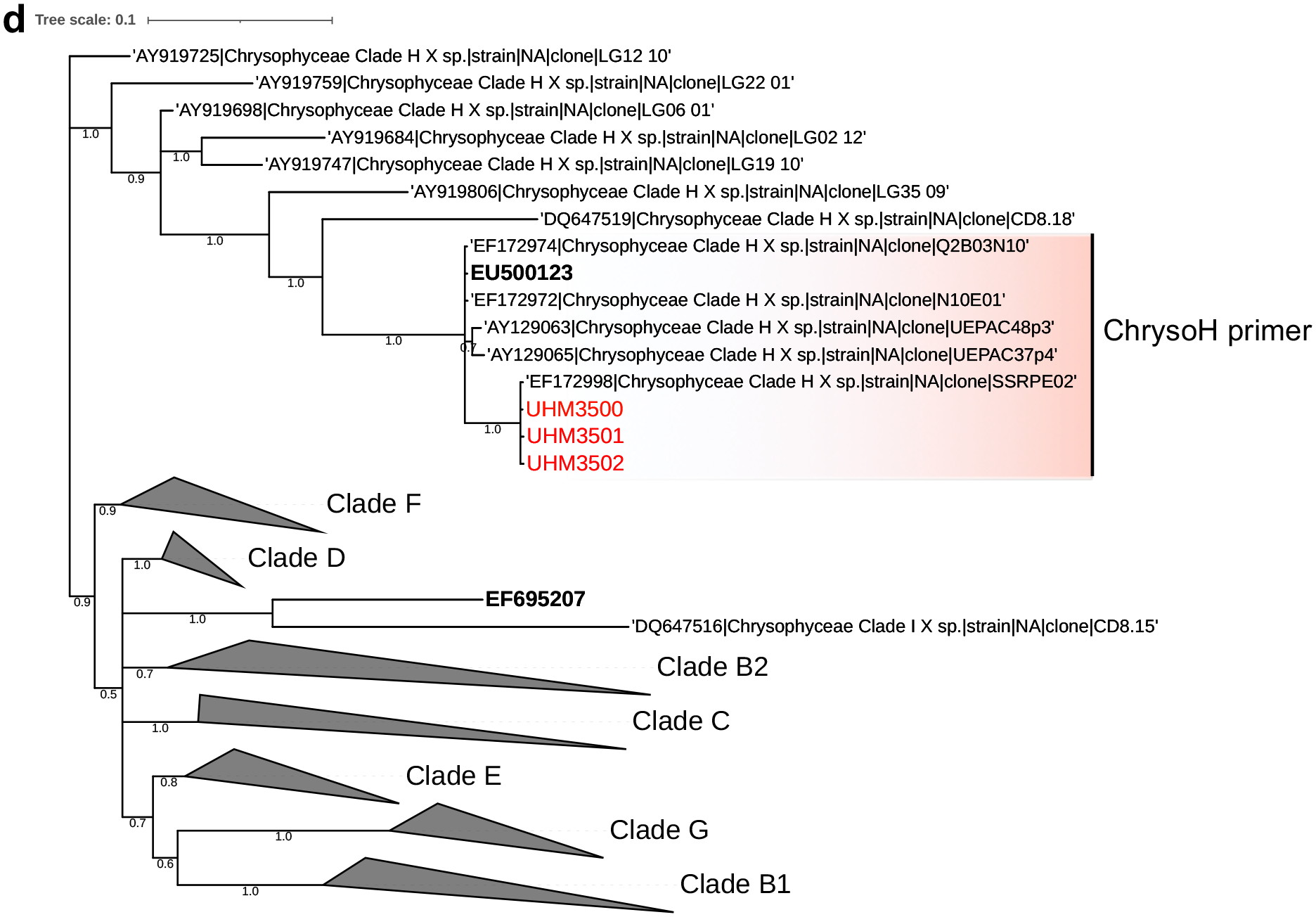
Phylogenetic analysis of the isolates used in this study. Bayesian majority consensus trees (with posterior probability shown on major branches) were inferred from partial 18S rDNA sequence alignments of our mixotroph isolates (in red), reference sequences from the PR^2^ database, and environmental sequences of putative grazers identified by Frias-Lopez et al. (2009) (in bold) for (**a**) Dictyochophyceae, (**b**) Haptophyceae, (**c**) Bolidophyceae and (**d**) Chrysophyceae. Six primer pairs theoretically target the majority, if not all grazers in the genera (or genus-level clades) *Florenciella*, DictyX, *Rhizochromulina*, Hap2, *Chrysochromulina* and ChrysoH. The *Triparma* primers target only some *Triparma* species, including *T*. *mediterranea* and *T*. *eleuthera*. The scale bar indicates the fraction of divergence.

**Supplementary Fig. S2.**
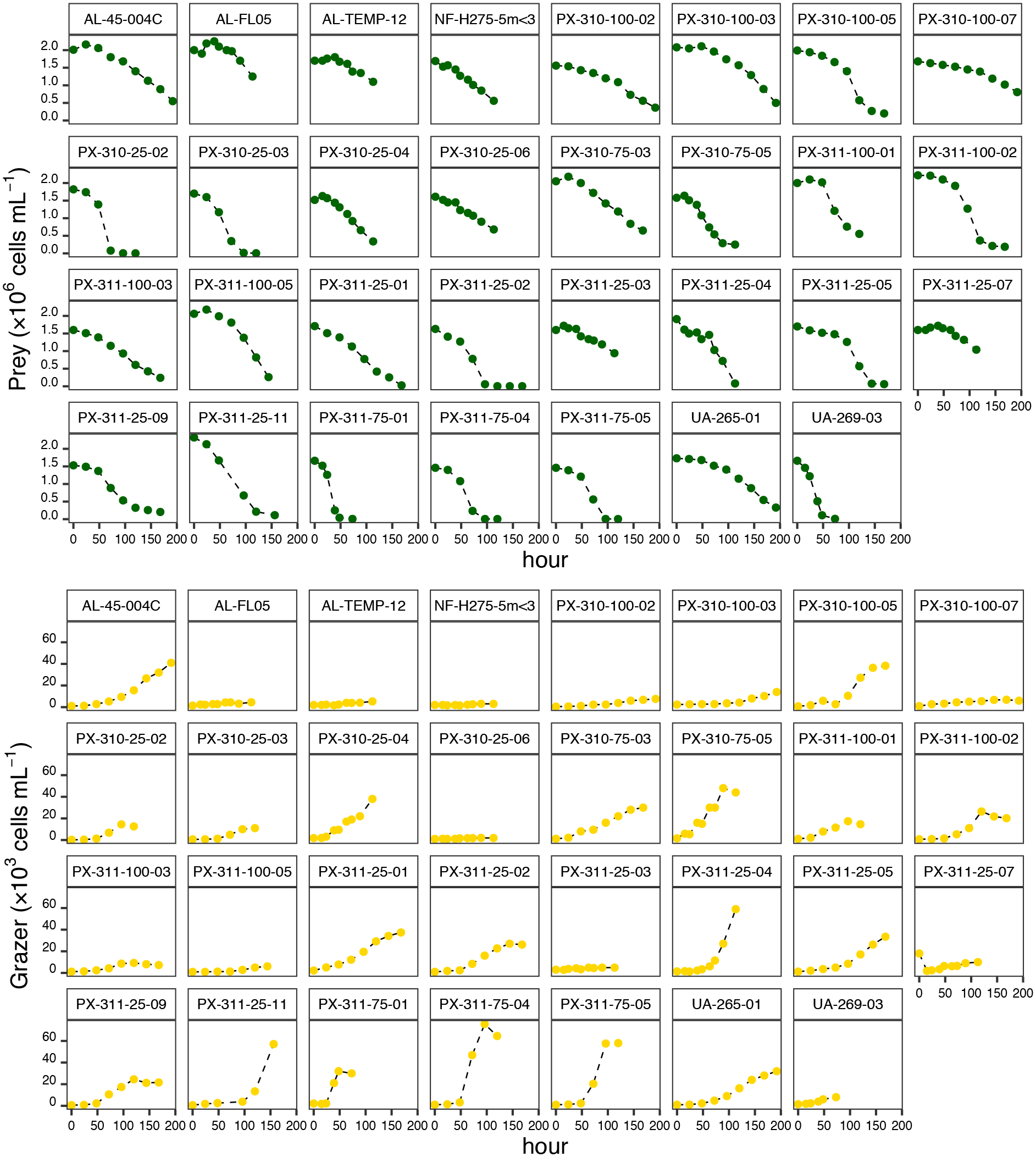
Changes in abundance of prey (upper panels) and grazer (lower panels) during grazing experiments. *Prochlorococcus* was the sole added prey and primary nitrogen source in the experiments. In the two types of controls (prey with no grazers or grazers with no prey) the prey and grazer abundances stayed roughly unchanged from day 1–5 at ∼2×10^6^ *Prochlorococcus* mL^-1^ or 100–1,000 grazers mL^-1^, respectively (data not shown). Replicated experiments for 11 isolates (Supplementary Table S1) have mostly yielded similar results and thus only data from one experiment each is shown here.

**Supplementary Fig. S3.**
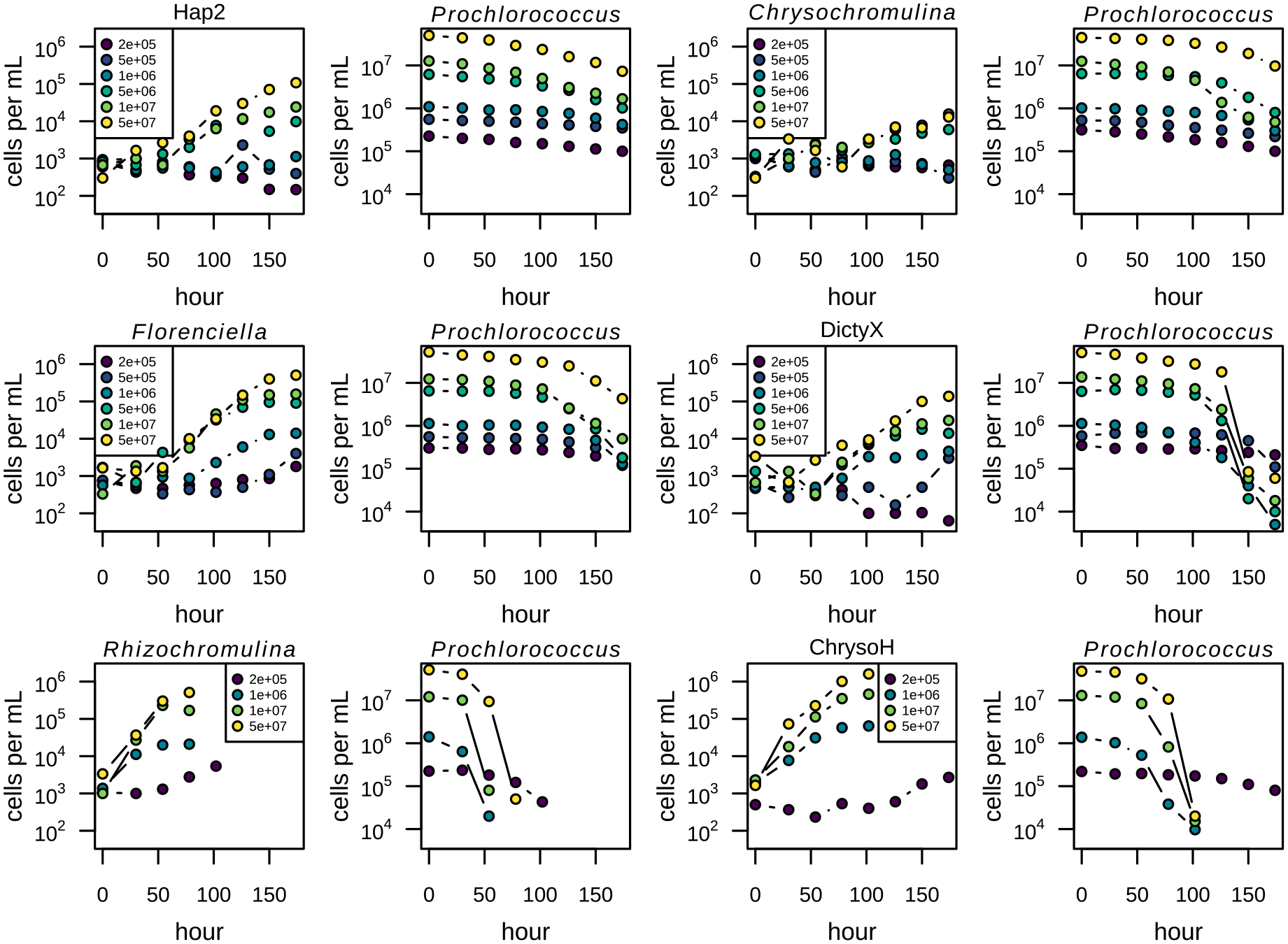
Grazer and prey trajectories (adjacent panels) of six isolates that were investigated for functional responses. Symbol colors indicate the concentration of added *Prochlorococcus* (cells mL^-1^) as shown in the legend.

**Supplementary Fig. S4.**
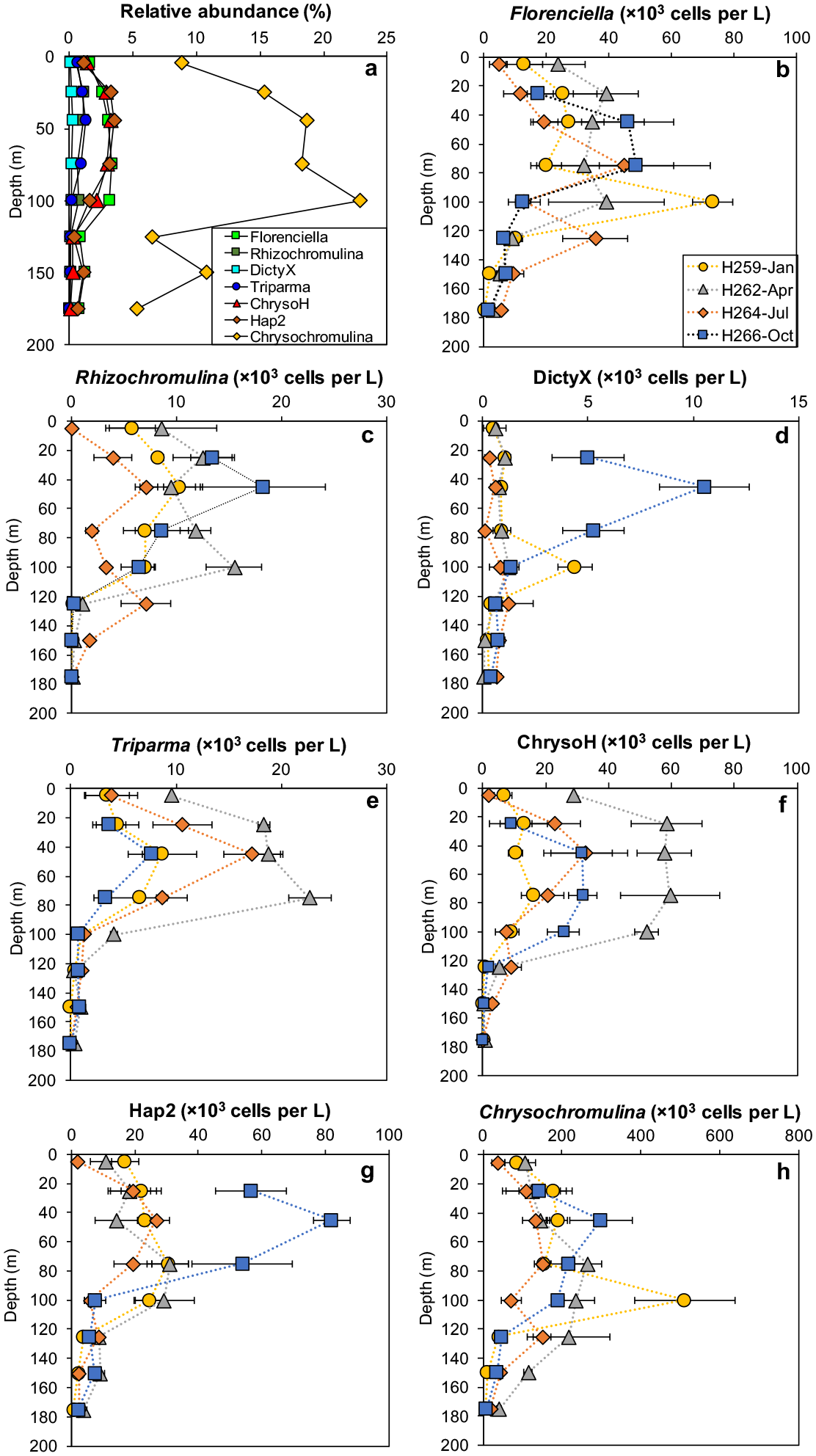
Relative and absolute abundances of seven mixotroph populations as a function of depth in the euphotic zone. (**a**) Annual average relative abundance as a function of depth, calculated by dividing the average absolute abundance of each group by the average flow cytometric counts of total pigmented eukaryotes over the same time period (data from at https://hahana.soest.hawaii.edu/hot/hot-dogs/). (**b**–**h**) Variations in the abundance of seven mixotrophic groups collected in four different seasons at Station ALOHA during Hawaii Ocean Time-series cruises #259, #262, #264 and #266, which took place in Jan, Apr, Jul and Oct of 2014, respectively.

**Supplementary Table S1.**
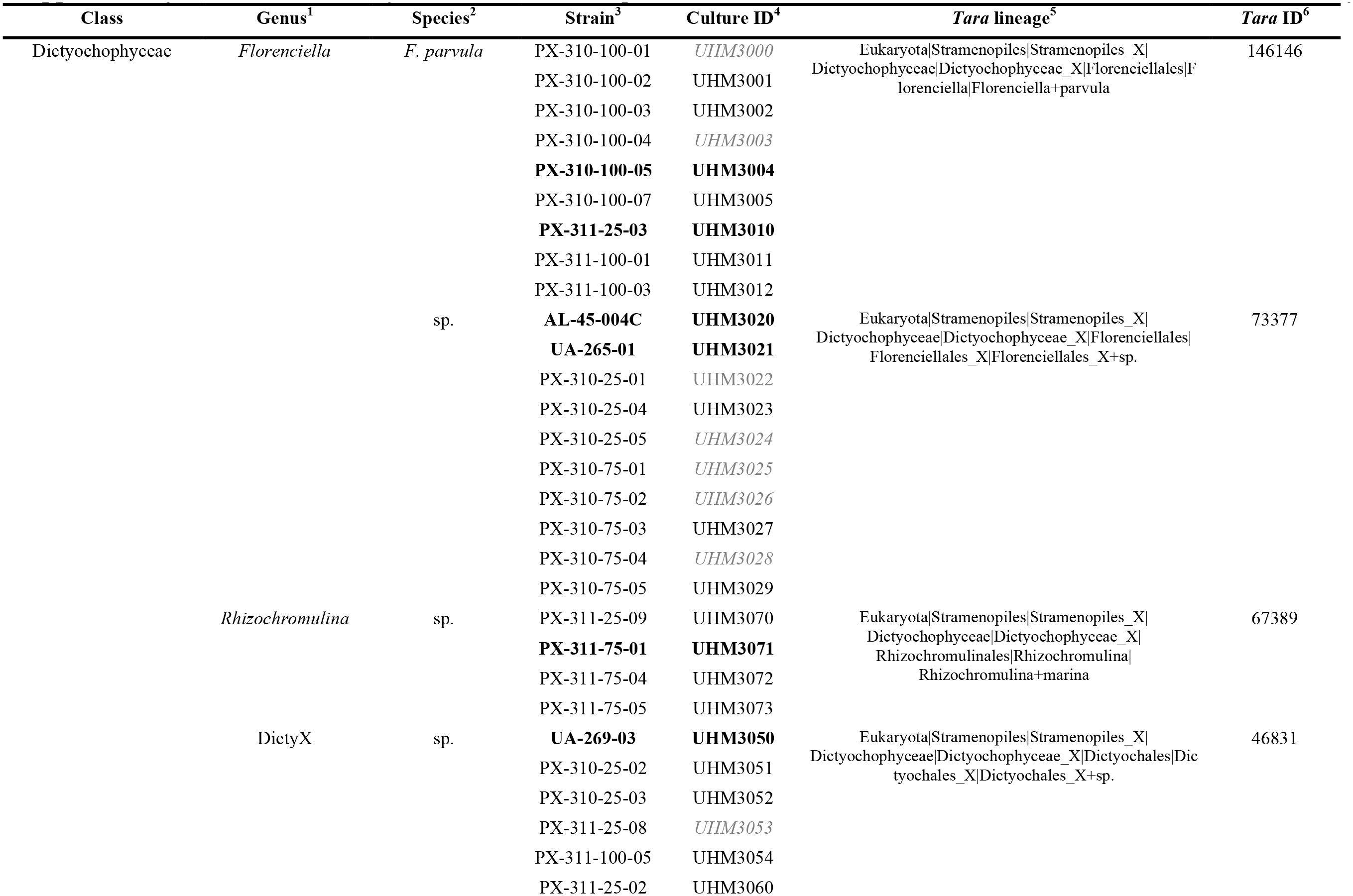
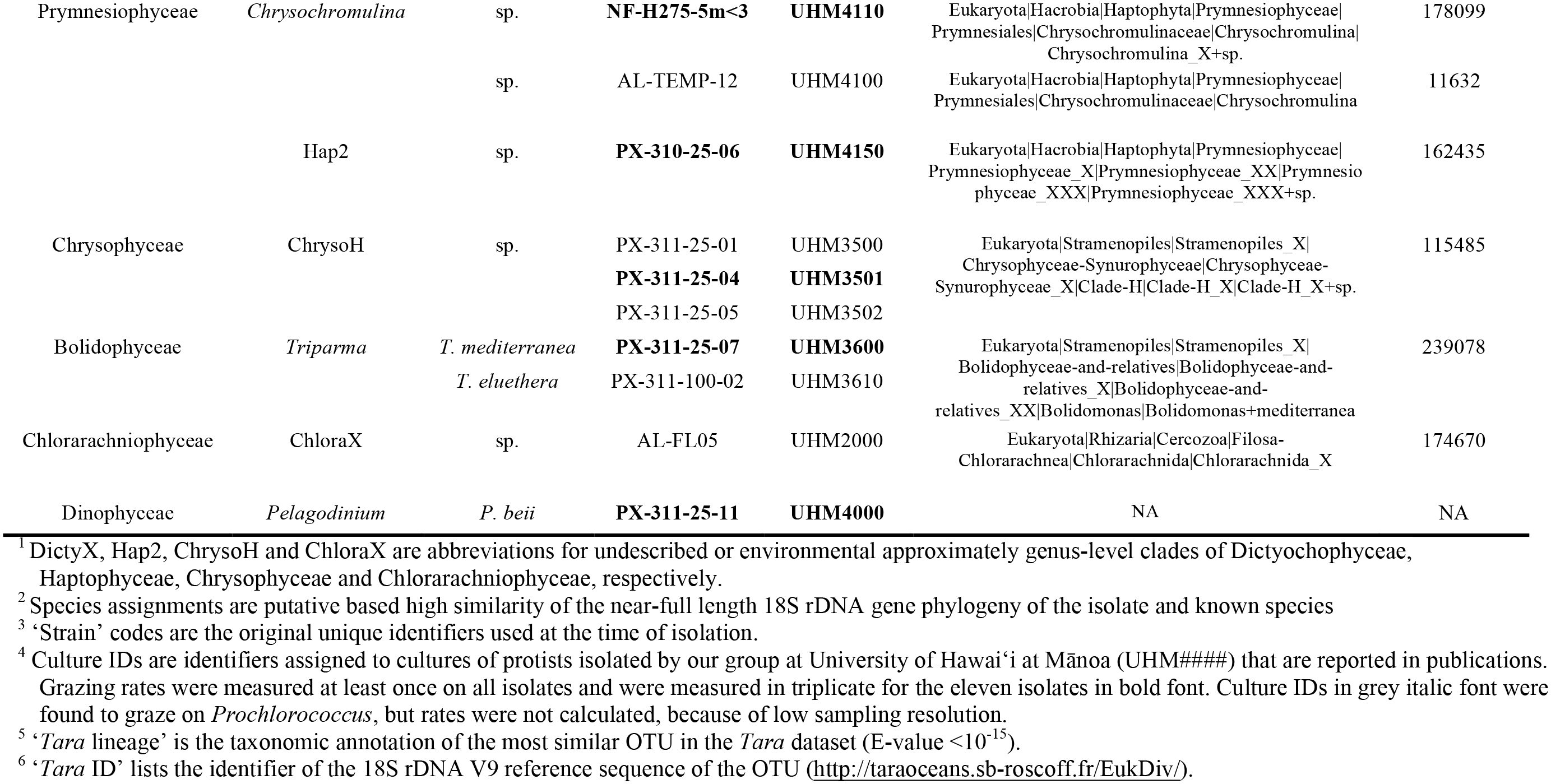
Taxonomy, culture IDs, and comparable *Tara* Oceans OTU information for isolates described in this study.

**Supplementary Table S2.**
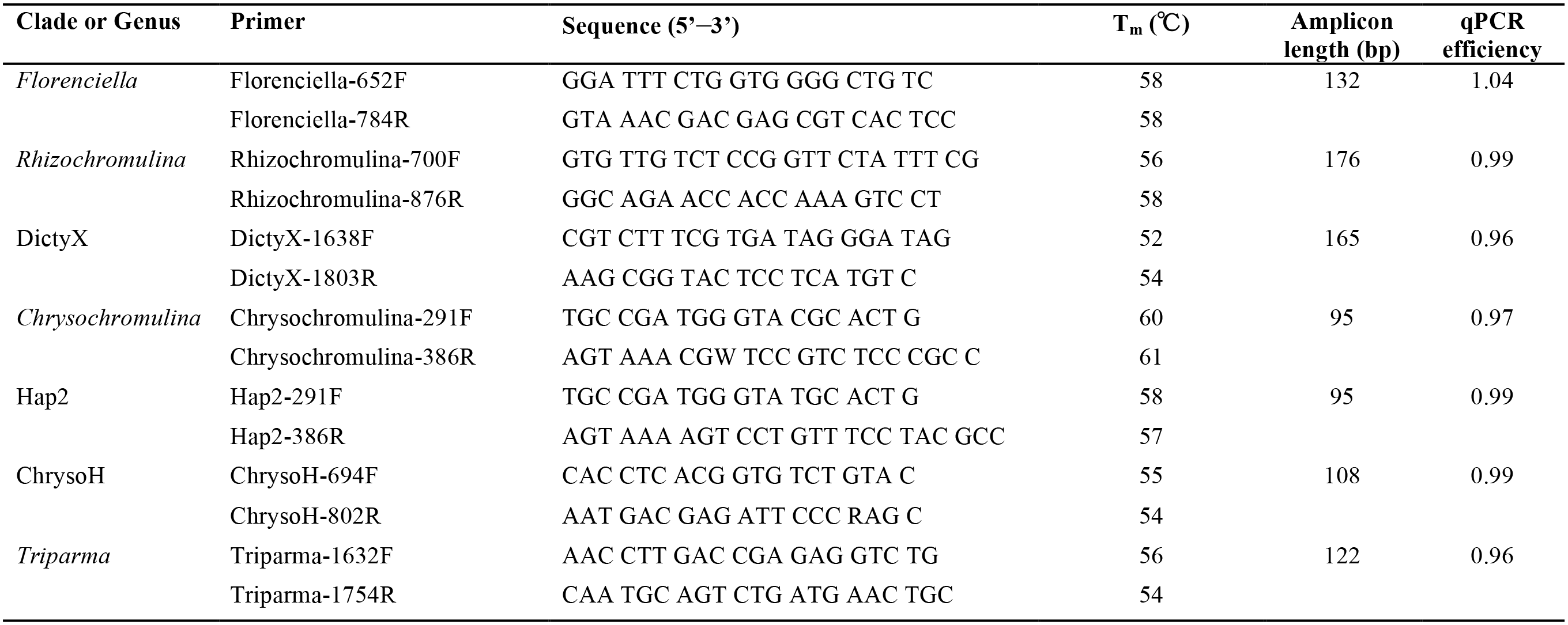
Detailed information of group-specific qPCR primers designed for targeting various mixotrophs in situ.

**Supplementary Table S3.**
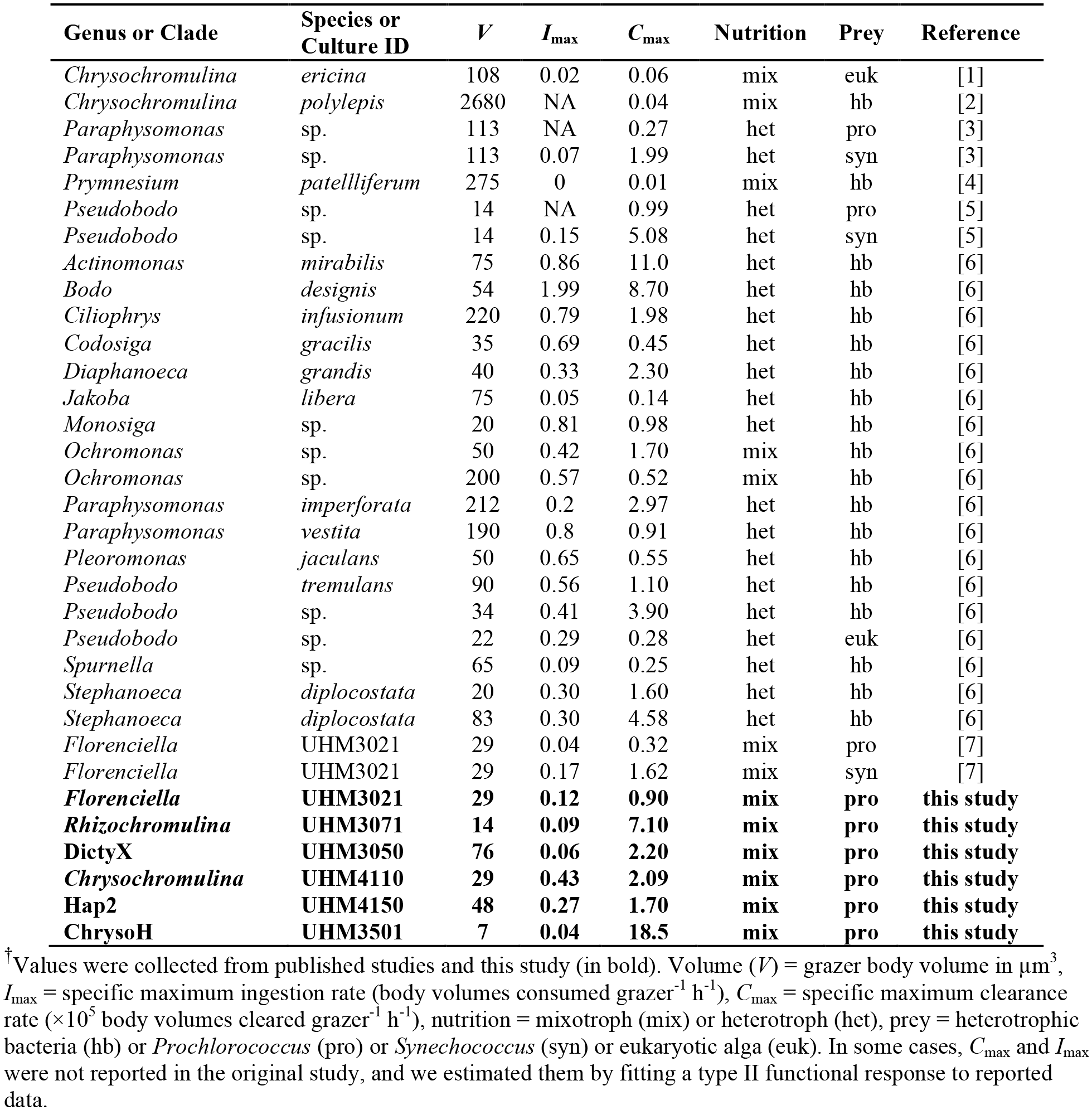
Maximum clearance rate and maximum ingestion rate parameters for heterotrophic and mixotrophic nanoflagellates^†^.

**Supplementary Table S4.** Nano-sized, non-dinoflagellate marine eukaryotes reported to have phago-mixotrophic capability^†^.

| Class | Genus | Size (µm) | Primary evidence of mixotrophy | Original publication |
| --- | --- | --- | --- | --- |
| Chlorarachniophyceae | *Cryptochlora* | >10 | ingestion of eukaryotic prey | [11] |
|  | *Lotharella* | >5 | presence of food vacuole | [12] |
|  | ChloraX | <5 | consumption and growth on *Prochlorococcus* | **this study** |
| Chrysophyceae | *Ochromonas* | >5 | ingestion of bacteria or FLB | e.g. [13][14] |
|  | *Dinobryon* | >3 | ingestion of latex-beads or FLB | [15][16] |
|  | ChrysoH | ~2 | consumption and growth on *Prochlorococcus* | **this study** |
| Cryptophyceae | *Geminigera* | >5 | ingestion of eukaryotic prey or microsphere | [17][18] |
|  | *Teleaulax* | >5 | ingestion of bacteria and *Synechococcus* sp. | [19] |
| Haptophyceae | *Chrysochromulina* | >3 | ingestion of bacteria or other prey | e.g. [20][21] **this study** |
|  | *Prymnesium* | >5 | ingestion of bacteria or other prey | e.g. [22][4] |
|  | *Haptolina* | >5 | ingestion of bacteria or other prey | [23][1] |
|  | Hap2 | ~5 | consumption and growth on *Prochlorococcus* | **this study** |
| Prasinophyceae | *Pyramimonas* | >10 | ingestion of FLB, microspheres or live bacteria | e.g. [24][18][25] |
|  | *Mantoniella* | <5 | ingestion of microspheres or FLB | [18][26] |
|  | *Cymbomonas* | >10 | ingestion of live bacteria | [27][25] |
|  | *Nephroselmis* | ~5 | ingestion of live bacteria | [9][25] |
|  | *Dolichomastix* | ~5 | ingestion of live bacteria | [25] |
| Raphidophyceae | *Heterosigma* | >10 | ingestion of *Synechococcus* | [28] |
| Dictyochophyceae | *Florenciella* | <5 | consumption and growth on *Prochlorococcus*, *Synechococcus* and bacteria | [7]; **this study** |
|  | *Rhizochromulina* | ~3 | consumption and growth on *Prochlorococcus* | **this study** |
|  | DictyX | 3-5 | consumption and growth on *Prochlorococcus* | **this study** |
| Bolidophyceae | *Triparma* | ~3 | consumption and growth on *Prochlorococcus* | **this study** |

<sup>†</sup>Taxa are mainly drawn from references [6][8][9] with original source references indicated. Specifically excluded are species that were isolated from benthic rocks or freshwaters, are larger than 20 µm, or for which evidence of bacterivory has been disputed (*Micromonas*, [10]).

## Notes

**Funding resource** This work was supported by National Science Foundation awards OCE 15-59356 and RII Track-2 FEC 1736030, and a Simons Foundation Investigator Award in Marine Microbial Ecology and Evolution (to K.F.E.).

### Competing Interest Statement

The authors have declared no competing interest.

